# Optimizing 3D-printed Scaffold Geometry Decreases Foreign Body Response and Enhances Allogeneic Islet Transplant Outcomes

**DOI:** 10.64898/2026.02.02.701816

**Authors:** Taylor R. Lansberry, Robert P. Accolla, Cameron C. Crouse, Irayme Labrada Miravet, Justin Walsh, R Damaris Molano, Camillo Ricordi, Cherie L. Stabler

## Abstract

Cellular therapy, such as beta cell transplantation for Type 1 diabetes, is a promising approach to durably alleviate disease states. Implanting cells within porous scaffolds is beneficial as they distribute the cells and mechanically support implantation; however, scaffolds can exacerbate foreign body responses (FBR). While the geometric features of a scaffold are known to impact FBR, there is limited consensus on what makes an ideal implant. Some have explored the role of pore size and interconnectivity; however, the impact of rung thickness between pores on FBR is broadly understudied. To investigate this parameter, we created a scaffold with reproducible geometric features and high biostability by combining 3D-printing with the polymer polydimethylsiloxane (PDMS). We tested 3D-printed scaffold prototypes with identical pore sizes but distinct PDMS rung thicknesses ranging from 150 to 300 µm. Upon transplantation, biocompatibility screening in a mouse model revealed that scaffolds with thicker PDMS rungs led to increased intra-device fibrosis. Additional spatio-proteomic analysis revealed distinct differences in host responses to rung changes, with alterations in macrophage and adaptive immune cell markers, as well as fibrotic proteins, within scaffolds containing thicker rungs. Selecting the optimized rung size, we evaluated its efficacy in rat syngeneic and allogeneic islet transplant models. In the allogeneic model, 3D-printed scaffold islet implants demonstrated robust efficacy and stability, yielding improved outcomes compared to PDMS scaffolds without optimized geometric features. Results from this study reveal how specific geometric scaffold features critically influence FBR to biomaterial implants, accelerating or mitigating fibrotic responses, and ultimately determining transplant success.

## Introduction

Cellular therapy involves the transplantation of donor cells to induce a therapeutic effect [1]. One prominent example is clinical islet transplantation (CIT) to treat people with Type 1 diabetes mellitus [2,3]. In this approach, donor islets are infused into the hepatic portal vein of the recipient to secrete insulin in response to changing blood glucose levels [4]. CIT has successfully restored physiological blood glucose control, yet few achieve long-term insulin independence [3]. This unfavorable long-term outcome is partially attributed to donor islet loss following poor engraftment within the intraportal site [5].

To alleviate this, researchers have explored the implantation of islets within devices at extrahepatic transplantation sites [6,7]. Device prototypes have varied from macroencapsulation approaches to highly porous scaffolds, with varying degrees of efficacy within preclinical models [8–11]. Several prototypes have been clinically screened, such as the Beta-O2 βAir, Viacyte PEC-Encap, and Viacyte PEC-Direct. The efficacy of these implants, however, has been modest. For example, the implantation of the Beta-O2 βAir, which encased islets within a macroencapsulation device supported by an oxygen tank, exhibited prompt fibrotic encapsulation post-transplantation, leading to a loss of detectable hormone release after only 5 days [12]. The Viacyte PEC-Encap, which housed islets in a planar device encapsulated within semi-permeable membranes, also failed due to aggressive fibrotic responses [13,14]. A redesign of the implant, whereby pores were drilled into the membrane to permit host cell infiltration into the device, improved c-peptide detection and decreased fibrous tissue, indicating that porous implants result in more favorable responses [15]. Overall, clinical outcomes to date highlight the deleterious impacts of foreign body responses (FBR) and implicate them as a key driver of cellular transplant failure.

Following the implantation of a device, host FBR is inevitable due to the trauma induced by the implantation procedure. While the timeline and degree of FBR to an implant can differ, common features include early non-specific protein adsorption, innate immune cell trafficking, macrophage polarization, foreign body giant cell formation, recruitment of fibroblasts, and eventually the formation of a collagenous avascular capsule [16–19]. More recent studies have also identified that adaptive immune cells play a role in this response; for example, T cells cross-talk with macrophages instigating foreign body giant cell formation [20–23]. While key cell players and pathways are still being uncovered, it is widely known that the degree of FBR to an implant varies depending on the implantation site, the cells it contains, and the material composition and geometric features of the device.

Several geometric features of an implant, including material surface texture, overall implant porosity, pore interconnectivity, and pore size, have been implicated in driving or diminishing FBR [24–27]. Current literature agrees that decreasing microscale surface roughness reduces inflammation and foreign body capsule formation [28–30], while increasing pore interconnectivity decreases collagen deposition and increases intra-device vascularization [31]. As such, porous scaffolds generally exhibit more favorable host responses than solid ones [32–34]. Regarding pore size, there is a lack of consensus on the ideal size range for limiting FBR, with some groups finding that smaller pores (<100 µm) induce macrophage polarization towards a pro-engraftment phenotype and decrease fibrous tissue ingrowth [32,35]. In contrast, others conclude that larger pore sizes (>400 µm) decrease foreign body capsule thickness [36,37]. One key geometric feature that may drive this variance, as it can impact both pore size and overall porosity, is rung thickness or the delta between individual pores [38–41]. Studies exploring the impacts of this scaffold feature are limited, likely due to challenges in controlling this parameter when using traditional fabrication techniques (e.g., particulate leaching, gas foaming, electrospinning) and/or degradable materials [7,42–45].

To investigate the role of rung thickness on FBR and graft efficacy, we combined a 3D-printing fabrication technique with a biostable polymer, polydimethylsiloxane (PDMS). PDMS is a flexible, non-degradable polymer with a robust clinical profile [43,46,47]. Using a reverse cast 3D-printing method, we fabricated 3D-printed PDMS scaffold prototypes with distinct geometric features. To investigate the impact of rung thickness on host responses, this parameter was varied from 150 to 300 µm, while the pore size was kept constant (300 µm). Prototypes were transplanted into immunocompetent mice, and host responses to these implants were assessed via Masson’s trichrome and immunohistochemistry staining. Additional spatial proteomic studies identified the distinct and local impact of the scaffold’s geometric features on host immune cell infiltration. After determining the optimal prototype, scaffolds were used to house pancreatic islets and implanted within diabetic syngeneic and allogeneic recipients to assess outcomes. Results found that scaffold features influenced host responses and overall therapeutic effects within our most challenging transplant model.

## Methods

### Fabrication of PDMS-based scaffolds

Scaffold fabrication was inspired by a previously published method for fabricating 3D-printed scaffolds using reverse casting [48,49]. Reverse-cast molds for fabricating our porous, 3D-printed scaffolds were modeled in Autodesk Inventor 2023 software and sliced for 3D-printing using Ultimaker Cura 5.2.1 software. Models were designed to fabricate three distinct scaffold geometries with a consistent intra-channel pore size of 300 x 300 x 100 µm (X, Y, Z), optimized previously [49], and interlocking PDMS rungs of different thicknesses in the X-Y dimension of 150 x 150 µm, 200 x 200 µm, or 300 x 300 µm and a consistent Z rung thickness of 100 µm (all 1.5 mm height). Other scaffold prototype geometries found in the supplementary material had a consistent modeled PDMS rung thickness of 150 x 150 µm (X-Y) and pore size of 300 x 300 (X-Y) with different rung and pore sizes of 100 µm and 200 µm in the Z dimension (all 1.5 mm height). Reverse-cast molds were 3D-printed on a commercially available 3D-printer (Model No. S3; Ultimaker) with a 0.4 mm nozzle (Ultimaker) using polyvinyl alcohol (PVA) filament (Cat. No. 8564; Amazon). Subsequently, a PDMS mixture consisting of a 4:1 (vol/vol) ratio of Part A to Part B (Cat. No. NC9017560; RTV 615; Momentive) was mixed in an automatic mixer (Model No. ARE-310; THINKY) until homogenous and added to the 3D-printed scaffold molds placed within a petri dish. Suspended gas within the PDMS solution was removed via a vacuum chamber for 1 hour. Scaffolds were then cured at 60°C overnight before removing excess PDMS from the top and bottom of the PVA molds. Molds embedded in cured PDMS were submerged in 500 mL of deionized water (per scaffold) for 72 hours, with the bath refreshed every 12 hours to fully dissolve the PVA mold. Finished PDMS scaffolds were dried overnight, and a 5 mm or 10 mm diameter biopsy punch was used to create experimental scaffolds for *in vivo* transplantation into mice or rats, respectively. Resulting scaffolds were stored under ambient conditions (20°C) until characterization or transplantation. Traditional scaffolds used for the allogeneic islet transplantation were fabricated using a salt-leaching technique as described previously [43].

### Characterization of Scaffold Structure

To assess the accuracy of the finished PDMS scaffold versus the theoretical model, the PDMS rung thickness and pore sizes were measured using brightfield microscopy and Fiji image analysis software [50]. Transmitted light imaging on a Leica TCS SP8 confocal microscope was used to image a population of PDMS rungs and pores in scaffold replicates before using Fiji to determine the average surface pore size and rung thickness (*N* = 4 – 6 scaffolds, *n* ≥ 40 rungs, *n* ≥ 40 pores). Average pore and rung sizes were compared to values provided by the modeling software; variance was measured and compared between distinct scaffold geometries to assess reproducibility. Due to the irregular pore sizes in the salt leached scaffolds, pore size was quantified by taking scanning electron microscopy (SEM) images of the scaffolds with image analysis.

Image analysis involved drawing two line segments across individual pores in the X and Y dimensions. The lengths of these line segments were input into equation (1) to calculate pore size for the salt-leached scaffolds.

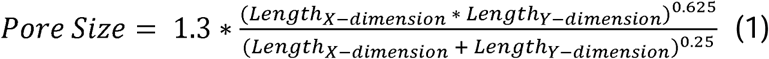

Macroscale bulk porosity was measured using a previously detailed mineral oil emersion method [51]. Briefly, scaffold height was measured using a digital caliper to calculate the cylindrical volume of the scaffold (*N* = 5 – 8). Scaffold dry weight was recorded before soaking the scaffold in mineral oil (Cat. No. 330779; Sigma Aldrich; density [ρ] = 0.838 g/mL at 25°C) and getting a wet weight. Porosity was calculated using equation (2). Porosity was compared to values provided by modeling software.

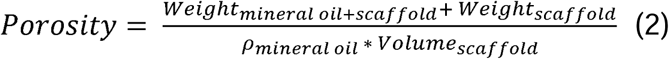

SEM was used to visualize the surface topography of both the 3D-printed and salt-leached scaffolds.

### Animal models

For biocompatibility studies, male *C57BL/6* mice (Jackson Laboratory) at 8 to 9 weeks old were used. Female *Lewis* rats (Envigo) at 180 to 200 grams were used as recipients for both syngeneic and allogeneic islet transplantations. Pancreatic islets were isolated from either healthy male *Lewis* or *Wistar Furth* rats weighing 250 to 280 grams. Animals were housed at either the University of Florida or the University of Miami under controlled environmental conditions in pairs with enrichment and free access to food and water. All animal protocols were approved by the University of Florida and the University of Miami Institutional Animal Care and Use Committee (IACUC) and were conducted following the NIH Guide for the Care and Use of Laboratory Animals, PHS Animal Welfare Policy, and the Animal Welfare Act.

### Evaluation of scaffold foreign body response and angiogenesis in C57BL/6 mice

Scaffolds (5 mm in diameter) with distinct PDMS rung thicknesses were sterilized via submersion in 70% isopropanol (Cat. No. A459; ThermoFisher) for 30 min followed by five 5-minute phosphate buffered saline (PBS; Cat. No. 10010023; Gibco) washes. To create a hydrophilic surface, scaffolds were incubated overnight in a 250 µg/mL human plasma fibronectin solution (Cat. No. 33016015; ThermoFisher) at 37°C. Scaffolds were then transferred to a custom-made, fitted PDMS mold for inclusion of a fibrin hydrogel within the empty pore spaces. Fibrin was fabricated by mixing fibrinogen and a master crosslinking enzyme solution in a 1:1 ratio. The fibrinogen solution consisted of fibrinogen (5.2 mg/mL; Cat. No. FIB3; Enzyme Research Laboratories), aprotinin (170 μg/mL; Cat No. 10981532001; Roche), and HEPES buffer (20 mM; Cat. No. 25–060-CI; Corning) containing sodium chloride (NaCl; 150 mM; Cat. No. S3014-500G; Sigma Aldrich). The master crosslinking enzyme solution consisted of calcium chloride (CaCl_2_; 9 mM; Cat. No. 449709-10G; Sigma Aldrich), thrombin (1.5 U/mL; Cat. No. T6884– 250UN; Sigma Aldrich), and HEPES buffer and was kept at 4°C until gelation. Excess fibronectin solution was aspirated and replaced with 10 µL of master crosslinking enzyme solution. Fibrinogen solution (10 µL) was immediately added to the master crosslinking enzyme solution inside the scaffold, mixed until homogenous, and allowed 1 min to crosslink.

For the surgery, male *C57BL/6* mice (Jackson Laboratory) were anesthetized using isoflurane (Cat. No. 50033; Covetrus) and a 3 cm midline ventral incision was made. The left and right lobes of the epididymal fat pad (EFP) were carefully exposed on sterile gauze outside of the body using sterile forceps. Both lobes were spread using sterile saline (Cat. No. 2B1324X; Baxter). One scaffold (5 mm diameter) containing the fibrin hydrogel was carefully placed onto each EFP lobe, wrapped by the edges of EFP, and sealed with a 15 µL fibrin hydrogel as detailed in a previous publication [52]. The study size was *n* ≥ 4 animals per scaffold geometry. Mice were monitored for changes in body weight and behavior for the duration of the study before euthanasia and scaffold explantation on day 35.

Scaffold grafts were explanted and fixed in 10% formalin for 24 hours before being cut in half to visualize cross-sectional areas. Then, scaffolds were histologically processed, paraffin-embedded, and sectioned. Histological staining included Masson’s trichrome stain (Cat. No. 87019; Epredia, Richard Allen Scientific) to visualize collagen deposition and immunohistochemistry to visualize vascularization.

Immunohistochemistry involved staining with anti-smooth muscle actin antibody conjugated with AlexaFluor 488 (Cat. No. AB184675; Abcam), anti-von Willebrand factor antibody (Cat. No. AB7356; Millipore Sigma), and DAPI (Cat. No. D1306; Invitrogen); a goat anti-rabbit antibody conjugated with AlexaFluor 568 was used as a secondary antibody to visualize the von Willibrand factor (Cat. No. A11036; Invitrogen).

For quantification of collagen deposition and nuclei inside the different 3D-printed scaffold pores, collagen and nuclei were segmented in the trichrome stained images using the Trainable Weka Segmentation plugin in Fiji ImageJ software (*N* ≥ 3 animals, *n* ≥ 2 ROIs per pore type in each image) [53]. Regions of interest (ROIs) of the major and minor pores were identified using the original trichrome images and added to the ROI manager for use on the segmented images. For the resulting segmented collagen images, the thresholds were adjusted to remove background and converted to binary. Then, the percent area of collagen coverage for each ROI was quantified. Alternatively, the segmented nuclei images were filtered using a Gaussian blur filter (Sigma = 0.8), thresholded until the background was removed, converted to binary, and watershed separation was applied. Using the analyze particles feature in Fiji, the number of nuclei between 3 – 50 µm in each ROI was quantified and normalized to the area of that ROI. Fold change in tissue compaction within the minor pores was calculated using equation (3).

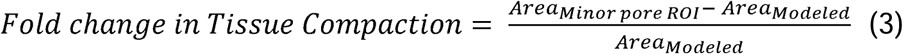

### Digital spatial profiling of scaffold explants

PDMS scaffolds with a geometry of 150 and 300 µm (X, Y) rung thickness were coated in fibronectin, filled with a fibrin hydrogel, and transplanted into the epididymal fat pad of *C57BL/6* mice as described above, before explanting and histologically processing the scaffolds on days 14 and 30. 7 µm sections of each scaffold type at the two timepoints were obtained from paraffin-embedded scaffold explant samples (*N* = 1; *n* ≥ 9 ROIs for each pore type, scaffold type, and time point). Samples were stained with a multiplexed cocktail of primary immune-centric antibodies tagged with unique UV-photocleavable oligonucleotides (30 proteins in total, Cat. No. 121300106 & 121300118; Nanostring Technologies; see **Supplemental Table S1**), rat anti-human CD3 antibody conjugated with AlexaFluor 647 (Cat. No. MCA1477A647; Bio-Rad Antibodies), mouse anti-rat α-Smooth Muscle Actin antibody conjugated with AlexaFluor 488 (Cat. No.

AB184675; Abcam), and Syto83 nucleic acid stain (Cat. No. S11364; Life Technologies).

The tissue was located using fluorescence imaging on the Nanostring GeoMx digital spatial profiling system and ROIs of each scaffold’s major and minor pores were selected (*n* ≥ 9 ROIs for each pore type within each scaffold/time point). ROIs were processed by focusing UV light through each ROI to release and collect the oligonucleotides from antibodies in each distinct area. After collection, the oligonucleotides for each ROI were hybridized to optical barcodes for digital counting using an nCounter Analysis System (Nanostring Technologies). Digital counts were normalized to the geometric mean of a positive internal spike in all ROIs to account for technical variability. Then, the counts were normalized to the geometric mean of housekeeping signals (i.e. histone H3 and S6) across all ROIs. The geometric mean of the normalized counts was compared between pore type, scaffold type, and explant time point to identify targets that were up- or down-regulated. To do this, the geometric mean of the normalized counts for one target was calculated across the ROIs for each of the comparators (i.e. major or minor pore). Then, the fold change was calculated by dividing the geometric mean of one comparator by the other for each target; this value was log_2_ transformed to look at proportional changes between the targets. These values were plotted on a volcano plot for visualization. Notably, if the geometric mean of a target count was below the geometric mean of the three background IgG antibody counts and not significantly up- or down-regulated for the comparison of interest, then the target was considered invalid.

### Islet isolation and islet loading into the scaffold

Donor pancreatic islets were isolated from healthy male *Lewis* or *Wistar Furth* rats (Envigo) via cannulation of the bile duct, injection of collagenase (Cat No. 5989132001; Roche), and removal of the pancreas. Then, the pancreas was enzymatically and mechanically digested before separating the islets from the tissue via a density gradient (Ficoll; Mediatech) as described in previous publications [43,54]. Islets were cultured at 37°C for 24 hours in CMRL 1066 media (Cat. No. 99-663-CV; Mediatech), supplemented with 10% fetal bovine serum (Cat. No. S1620; Biowest), 1% penicillin-streptomycin (Cat. No. 30-002-CI; Corning), and 1% L-glutamine (Cat. No. 25-005-CI; Corning). Then, the islets were counted using an algorithm for calculating 150 µm diameter islet equivalent (IEQ) number before loading the islets into the scaffolds [55].

As described above, 10 mm diameter 3D-printed and salt-leached scaffolds were sterilized and coated in fibronectin for 24 hours. A fibrinogen solution and master crosslinking enzyme solution were fabricated, as also described. Islet aliquots containing 2,000 IEQ *Lewis* islets (for syngeneic transplant) or 3,000 IEQ *Wistar Furth* islets (for allogeneic transplant) were loaded into the scaffold by concentrating each aliquot into 39 µL of the fibrinogen solution. 10 mm diameter 3D-printed scaffolds were transferred to a custom-made, fitted PDMS mold, and excess fibronectin solution was removed. 9 µL of the master crosslinking enzyme solution was added to each scaffold within the fitted mold. Next, the 39 µL solution of fibrinogen containing islets was carefully added inside each scaffold. Finally, the remaining 30 µL of master crosslinking enzyme solution was added to the scaffold to complete the 1:1 fibrin hydrogel ratio. The hydrogel was allowed 1 min to crosslink and seal the islets inside the 3D-printed scaffold. Islet loading into the 10 mm diameter salt-leached scaffolds was performed as described in a previous publication [43].

### Diabetes induction and evaluation of islet transplant outcomes

*Lewis* rats were rendered diabetic via two intraperitoneal injections of streptozotocin (Dose 1: 55 mg/kg, Dose 2: 50 mg/kg, 3 days apart; Cat. No. S-0130; Sigma-Aldrich), a β-cell toxin. Rats were only used as scaffold recipients if they exhibited three consecutive daily non-fasting blood glucose readings > 350 mg/dL via a portable glucometer (One Touch Ultra 2, Model No. 353885011853; Lifescan).

Recipient animals in the allogeneic islet transplant were immunosuppressed using a three-drug regimen. Anti-lymphocyte serum (ALS, Cat. No: AIA5940/20; Accurate) was administered in two intraperitoneal injections pre-operative day 1 and post-operative day (POD) 2 (dosage determined for each lot number of ALS via a titration study). Mycophenolic acid (MPA; Cat. No. 0078-0385-66; Novartis) was administered daily following scaffold-islet transplantation via oral gavage at 20 mg/kg until POD 14. At POD 15 and 16, the dose was lowered to 15 mg/kg, POD 17 and 18 to 10 mg/kg, and POD 19 and 20 to 5 mg/kg before completely removing MPA after POD 20. Fingolimod (FTY-720; Cat No. orb154632; Biorbyt) was administered daily via oral gavage at a concentration of 1 mg/kg for the entire duration of the study.

For both syngeneic and allogeneic islet transplants, under general anesthesia using isoflurane, a 5 cm midline incision was made on the rat’s ventral side. The omental pouch was exposed from the peritoneal cavity and spread onto sterile gauze using sterile saline. An individual islet-loaded scaffold (*N* = 4 animals per scaffold group in syngeneic transplant; *N* = 7 animals per scaffold group in allogeneic transplant) was placed on the omentum before wrapping the edges of the omentum around the scaffold and sealing with a 30 µL fibrin hydrogel. The omental pouch was placed back inside the peritoneal cavity, and the incision was sutured and stapled closed. Blood glucose and weight of the recipients were measured daily. Functional scaffold-islet grafts were defined as animals having a stable blood glucose reading < 200 mg/dL for three consecutive days.

For the syngeneic study, grafts were assessed for functionality using an intravenous glucose tolerance test on POD 34 for the 3D-printed scaffold group and POD 26 for the salt-leached scaffold group. Animals were fasted overnight and anesthetized via isoflurane before injecting a 50% dextrose solution (Cat No. C1880150; Covetrus) intravenously at a concentration of 2 µL per gram of body weight. Blood glucose was quantified periodically over 2 hours. All scaffolds were explanted after POD 30 and processed for histological analysis.

For the allogeneic study, time to normoglycemia was determined by recording the first POD that each animal had a stable blood glucose reading after three consecutive days of stable readings. Scaffolds were explanted as each animal rejected their graft and had three consecutive blood glucose readings > 300 mg/dL. POD to reject the graft was determined by recording the first POD that each animal had an unstable blood glucose reading > 300 mg/dL if the animal had three consecutive unstable daily blood glucose readings. The remaining animals at POD 92 (3D-printed scaffolds) and POD 105 (salt-leached scaffolds) were sacrificed, and their scaffold grafts were explanted. All scaffold explants were fixed in 10% formalin buffer and processed for histological analysis. Scaffold and tissue sections were stained with Masson’s trichrome staining and hematoxylin and eosin to view collagen, vascularization, and islet integration.

### Statistical analysis

Statistical differences were analyzed using Graphpad Prism 10 software, and results were expressed as the mean ± SD. Depending on the group and parameter sizes, different tests were employed. For equal variance (passed F test), two experimental groups were analyzed using a Student’s T test, > 2 groups via one-way ANOVA with Tukey’s multiple comparisons test, or more than two parameters via two-way ANOVA with Sidak or Tukey post hoc analysis. For unequal variances, a Brown-Forsythe ANOVA test with a Dunnett’s T3 multiple comparisons test was used instead. For digital spatial profiling, immune cell targets were presented as fold changes in volcano plots. Comparisons between explant day and scaffold type were analyzed using a Linear Mixed Model with Benjamini-Hochberg multiple test correction in Nanostring GeoMx DSP Analysis Suite Version 3.1.0.222 to account for the idiosyncratic biologic variability between two different mouse samples. Comparisons between pore type within the same mouse sample were analyzed using multiple unpaired T-tests. For islet transplant efficacy studies, the time to normoglycemia and rejection rates were analyzed using a Mantel-Cox (log rank) test to compare the 3D-printed scaffold to the salt-leached scaffold. The number of intra (*n*) or inter (*N*) experimental replicates and the specific statistical method conducted for each study are reported in the figure legends. Details on the statistical methods and results are summarized in **Supplementary Table S2**. Differences for all experiments were considered significant if *p < 0.05*.

## Results

### Geometric Fidelity and Structural Properties of 3D-Printed and Salt-leached Scaffolds

To assess the impact of distinct geometric scaffold features on FBR and islet transplant outcomes, we fabricated scaffolds using a 3D-printing technique. This method can achieve high control over geometric features and length scales. Additionally, the scaffold was generated using biostable PDMS, which is expected not to degrade *in vivo* during the implant period. To fabricate these scaffolds, reverse-cast, sacrificial molds were 3D-printed using water-soluble polyvinyl alcohol (PVA) before integrating uncured PDMS into the molds (**Figure 1A**). After degassing and curing, the sacrificial mold was removed via hydration, leaving a flexible PDMS scaffold with distinct geometries.

**Figure 1:**
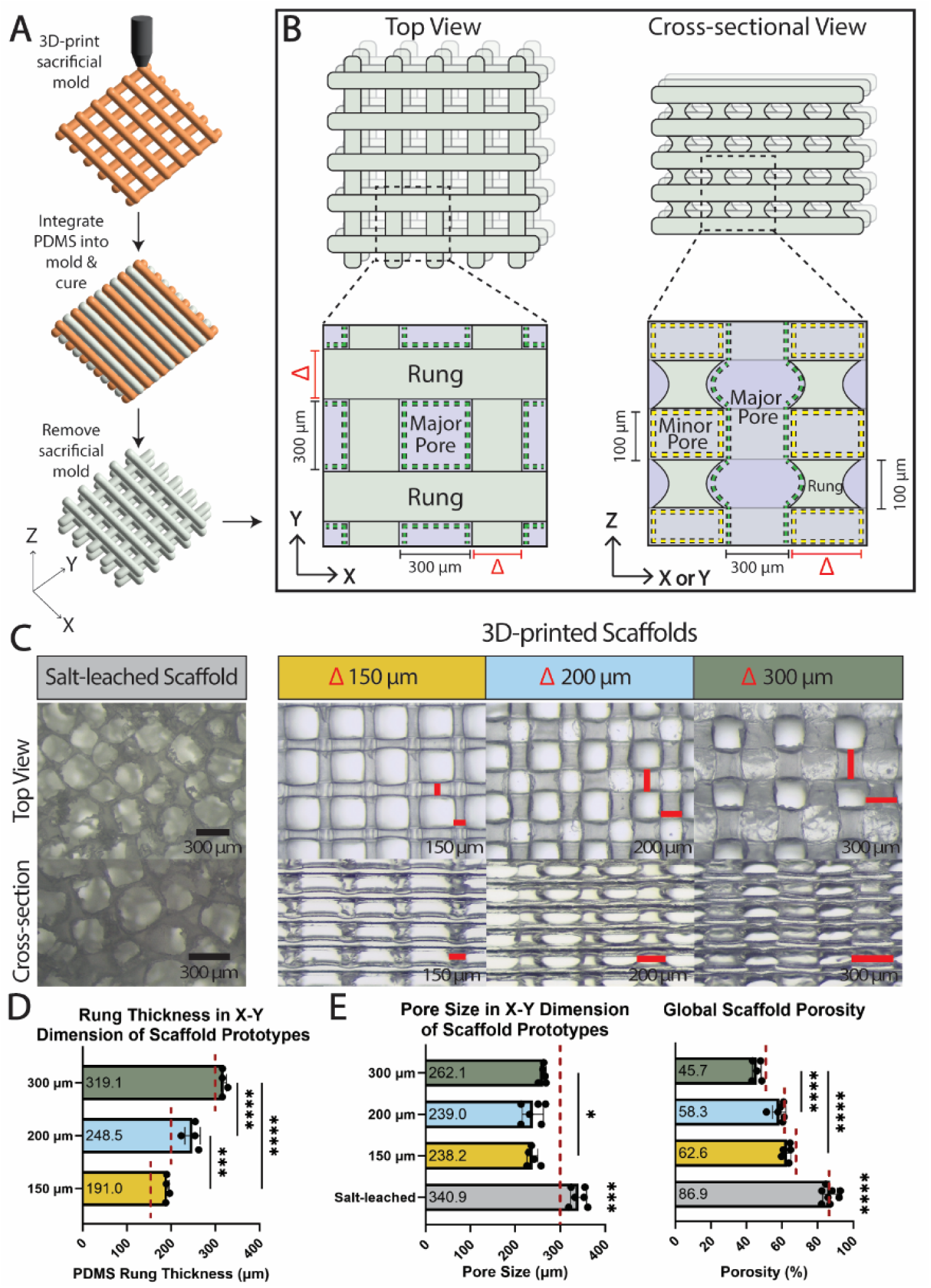
Fabrication and evaluation of 3D-printed scaffold prototypes with distinct PDMS rung thicknesses. Reverse cast method for 3D-printing the scaffold prototypes (A). Top and cross-sectional views of the scaffold prototypes. Scaffold geometry was altered by modulating rung thickness in the X-Y dimension (red Δ). A rung thickness of 100 µm in the Z dimension and pore size of 300 x 300 x 100 µm in the X-Y-Z dimension was kept consistent. Modulating rung thickness inherently altered major (green dashed line) and minor (yellow dashed line) pore sizes (B). Top and cross-sectional brightfield images of a traditional salt-leached scaffold in comparison to the 3D-printed scaffold prototypes with 150, 200, or 300 µm rung thickness in the X-Y dimensions (C). Quantification of rung thickness in the X-Y dimension of 3D-printed scaffold prototypes (D). Quantification of pore size in the X-Y dimension and global porosity of 3D-printed and salt-leached scaffolds (E). Theoretically derived values for each parameter (red dotted line) are shown for reference. (*N* = 4 – 6 scaffolds used for all measurements, *n* ≥ 40 rungs or pores per scaffold; data represented as mean ± SD; One-way ANOVA with Tukey’s multiple comparisons test for rung thickness and porosity measurements, Brown-Forsythe ANOVA test with Dunnett’s T3 multiple comparisons test for pore size measurements; * *p* < 0.05, *** *p* < 0.001, **** *p* < 0.0001).

The geometric features of the 3D-printed scaffold could be manipulated across all three X-Y-Z dimensions (**Figure 1B**). From the top view, geometric parameters that could be altered included the pore size within the X-Y plane (designated as the ‘major pores’, green dashed line) and rung thickness (i.e., the space between major pores). Along the cross-sectional plane, the geometry of the pores that connected the major pores (designated as the ‘minor pores’, yellow dashed line) was dictated by both the rung thickness (along the X or Y direction) and the distance between the 3D-printed layers (along the Z direction).

We previously identified that a major pore size of 300 x 300 µm^2^ along the X-Y plane supported robust host cell integration and vascularization [49]. With this parameter fixed, we next set out to investigate the impact, if any, of rung thickness (denoted as Δ in **Figure 1B**). Prototypes with a rung thickness between 150 to 300 µm, but consistent 300 x 300 x 100 µm X-Y-Z major pores, are shown in **Figure 1C**. Examining both top and cross-sectional views of the resulting PDMS scaffolds, all 3D-printed prototypes exhibited consistent pore sizes and pore connectivity with smooth rungs, in stark comparison to PDMS scaffolds fabricated using a traditional compression and salt-leaching method, as shown via SEM imaging (**Supplementary Figure S1**).

To validate the fidelity of the prototypes, scaffold rung thickness and pore size were quantified using image analysis. As expected, PDMS rung thickness significantly differed across the three scaffold prototypes (**Figure 1D**, *p < 0.0001*); however, experimental measurements were slightly larger than their modeled rung thickness (**Figure 1D**, red dashed line) with experimental values 5.3 ± 5.3%, 24.3 ± 8.8%, and 27.4 ± 2.7% higher than their modeled size for the 300, 200, and 150 µm rung scaffolds, respectively. This increased rung thickness resulted in a slight decrease in the average major pore size for the scaffolds, with reductions of 12.6 ± 2.3%, 20.4 ± 7.8%, and 20.6 ± 4.1% for the 300, 200, and 150 µm rung scaffolds, respectively (**Figure 1E**). These results indicate a trend of decreased printing accuracy with decreased rung thickness, as shown by the percent change in pore and rung size from their modeled dimensions (**Supplementary Figure S2,** *p = 0.0008 for rung thickness*; *p = 0.0589 for pore size,* ). Comparing the pore size of the 3D-printed scaffolds to those fabricated using a traditional salt-leaching technique found pore sizes to be of similar scale (∼300 µm) but significantly different (**Figure 1E***, p < 0.0001*); the intra-scaffold variability of pore sizes within salt-leached scaffolds were significantly more variable than the 3D-printed scaffolds as summarized in the pore size variability plot (**Supplementary Figure S3**, *p < 0.0001*). Global porosity measurements capture the total open space within each scaffold type. CAD models predicted a decrease in overall porosity correlating with increasing rung thickness; experimental measurements generally matched model predictions (**Figure 1E**, *p = 0.3435)*.

Scaling up the scaffold to a size more relevant for clinical translation can be challenging. For traditional fabrication methods that require the compression of particulate/polymer mixtures during the curing process to ensure high pore interconnectivity, such as the salt-leaching method employed here, increasing the scaffold diameter typically results in the need for complex equipment to deliver the high compression requirements for larger surfaces [45,56]. Alternatively, 3D-printing methods do not have this limitation [57]. To evaluate the capacity to scale our reverse casting 3D-printing method, 80 mm diameter PDMS scaffolds were fabricated. The resulting scaffolds had geometric features that were consistent with their 43-fold smaller counterparts with high fidelity and low intra-scaffold variability, as shown in the rung and pore size plots (**Supplementary Figure S4 and Video S1,** *p = 0.0804 for rung thickness, p = 0.7221 for pore size*).

### Evaluation of Foreign Body Response to 3D-Printed Scaffold Prototypes

Following the validation of prototype fabrication, we next sought to understand how rung thickness impacts host tissue integration and overall compatibility.

Immunocompetent *C57BL/6* mice were used for this screen, as they are considered a “fibrosis-prone” strain that provides a robust screening of material compatibility [58–61]. All three 3D-printed scaffold prototypes were tested via implantation within the EFP, which is a common mouse extrahepatic site for islet transplantation (**Figure 2A**) [25,49,62]. Scaffolds were explanted after 30 days to assess foreign body responses. Visual inspection of histological sections after 30 days *in vivo* identified distinct spatial differences in collagen deposition across the different prototypes. Within the major pores of the scaffolds (**Figure 2B**, green boxes), host integration and extracellular matrix (ECM) deposition, as classified by Masson’s trichrome, appeared similar across all scaffolds; however, a distinct elevation in collagen within the minor pores was visualized within prototypes with rung thickness ≥ 200 µm (**Figure 2B**, yellow boxes). For the largest rung thickness of 300 µm, this fibrotic region often expanded into the major pore space.

**Figure 2:**
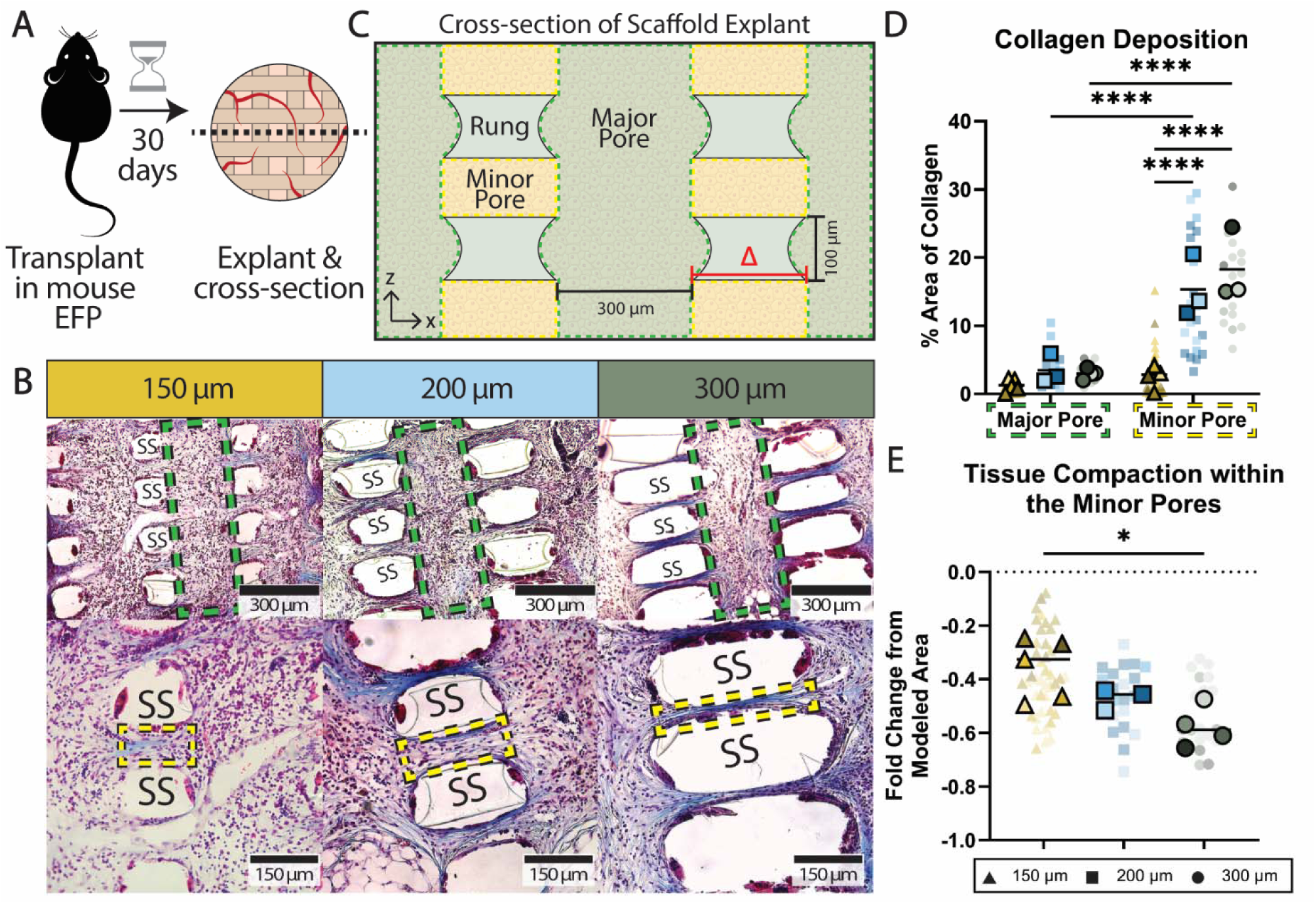
Imaging and quantification of host tissue integration into scaffold prototypes. Scaffolds were transplanted individually into the *C57BL/6* mouse epididymal fat pad, explanted, and cross-sectioned after 30 days (A). Cross-sectional images of Masson’s trichrome stained explants at 10x (top row) and 20x (bottom row) magnification (B). Schematic depicting the segmentation of the major (green) and minor (yellow) pores within the cross-sectioned explants (C; red Δ -change in scaffold rung thickness). Quantification of collagen deposition within the major and minor pores of the scaffold explants (D). Quantification of tissue compaction within the minor pores of the scaffold explants (E). (SS = silicone scaffold rung, green dashed boxes – major pore, yellow dashed boxes – minor pore; *N* = 3 – 5 scaffold explants, *n* ≥ 3 minor pore ROIs per scaffold, *n* ≥ 2 major pore ROIs per scaffold, shade of color indicates different mouse sample; all data points with mean shown; Two-way ANOVA with Tukey’s multiple comparisons test for collagen quantification, One-way ANOVA with Tukey’s multiple comparison’s test for compaction quantification; * *p* < 0.05, **** *p* < 0.0001).

These unexpected spatial differences were analyzed via a segmented image analysis (**Figure 2C**). Globally, pore type and scaffold geometry significantly impacted intra-device collagen deposition (**Figure 2D**, *p < 0.0001*). While no significant differences were detected across scaffold prototypes within their major pores, significantly more collagen was quantified within the minor pores of the 300 and 200 µm scaffolds, when compared to the 150 µm scaffold (18.28 ± and 15.37 ± 4.56% vs. 2.84 ± 1.63%, *p < 0.0001*). Additionally, within the 300 and 200 µm scaffolds, significantly more collagen was found within their minor pores compared to their major pores (*p < 0.0001*); this was not the case for the 150 µm scaffold (*p = 0.3760*). Collagen deposition appeared to visually impact tissue compaction within the minor pore areas, particularly when comparing the 150 and 300 µm scaffolds. Statistical analysis found that as rung thickness increased, host tissue within the minor pores was more compact (**Figure 2E**; *p = 0.0178*). Quantifying cell density, no significant differences were detected between pore or scaffold types (**Supplementary Figure S5,** *p = 0.3519 for pore type, p = 0.8505 for scaffold type*). Intra-device vascularization of these explants was assessed via immunohistochemistry co-staining of von Willibrand factor and α-smooth muscle actin (SMA); these images are summarized in **Supplementary Figure S6.**

To explore if the height of the minor pores (i.e. the Z-direction) would have an impact, additional prototypes were generated in which the Z height of the PDMS rungs was varied from 100 to 200 µm while the major pore size (300 x 300 µm^2^) and rung thickness (150 µm) in the X-Y dimension remained constant; fidelity data for these prototypes are summarized in **Supplementary Figure S7**. Biocompatibility studies of these scaffold prototypes did not detect any significant impacts of minor pore height on collagen deposition (*p = 0.6722*; two-way ANOVA). Cell density was globally significant between scaffold types (*p = 0.0121*; two-way ANOVA), but post-hoc analysis did not reveal any significant differences between any particular groups (**Supplementary Figure S8**).

### Quantitative Proteomic Digital Spatial Profiling of Host Responses to 3D-printed Scaffold Prototypes

To identify the immune cell phenotypes that drove the stark histological differences in fibrotic tissue ingrowth within these 3D-printed scaffolds, proteomic digital spatial profiling was employed. For this study, scaffolds with either 150 or 300 µm rung thickness were implanted into the EFP of *C57BL/6* mice. Scaffolds were then explanted on days 14 and 30 to assess host responses at early-(day 14) and advanced-stage (day 30) FBR [63]. Histological sections were stained to identify ROIs, and immune-relevant antibodies conjugated with UV-photocleavable oligos (25 in total) were added (**Figure 3A**); the immune-relevant antibodies used are listed in **Supplementary Table S1.** ROIs were identified within either major (green-white dotted boxes) or minor (yellow-orange dotted boxes) pores for each scaffold (**Figure 3B**). The levels of immune-related proteins per ROI were calculated and normalized to the housekeeping proteins within their respective ROI.

**Figure 3:**
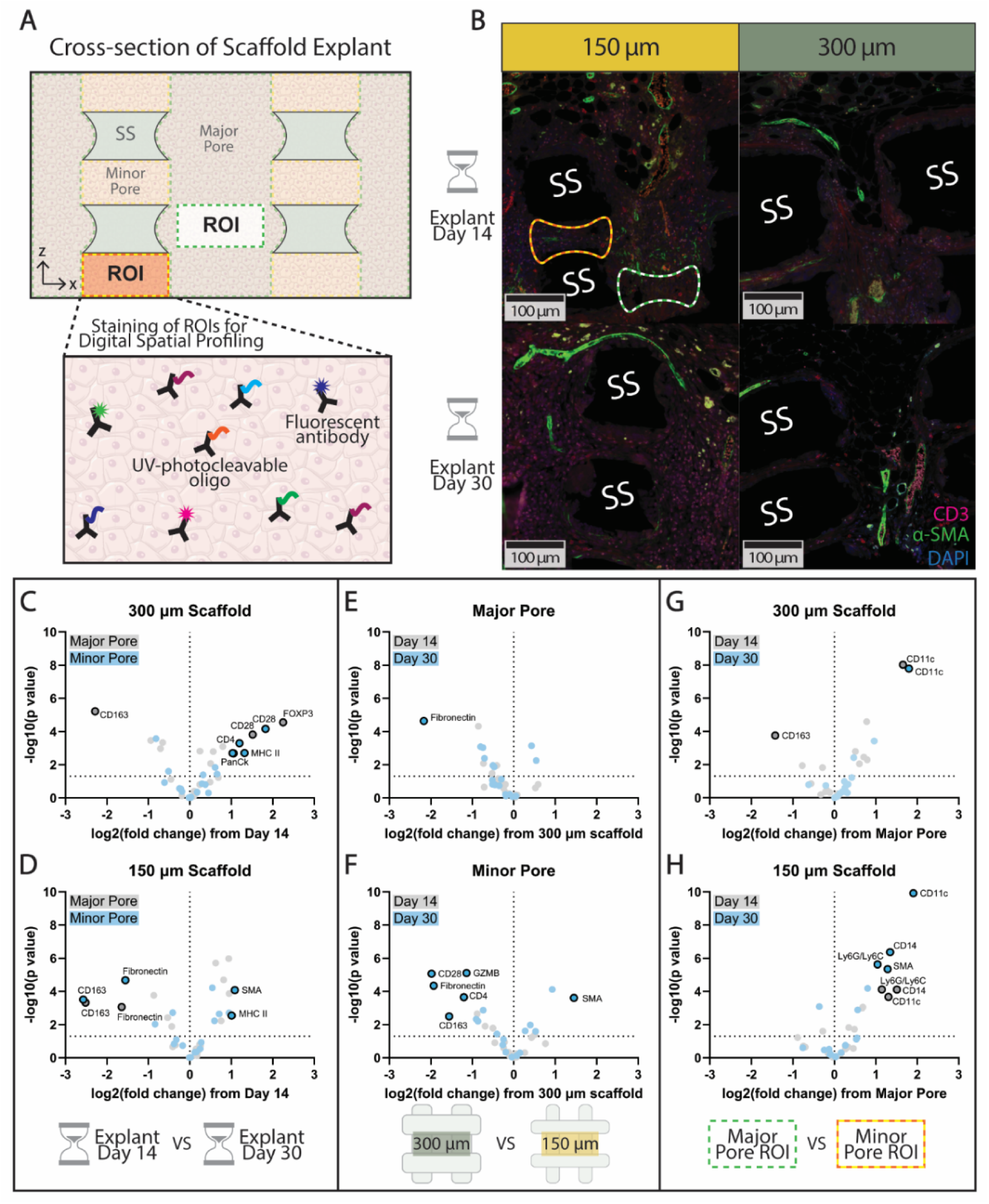
Digital spatial profiling of 150 µm and 300 µm 3D-printed scaffold prototypes explants after transplantation into the epididymal fat pad of *C57BL/6* mice for 14 and 30 days. Schematic depicting the segmentation of cross-sectioned explants into major and minor pore ROIs, cutout depicts staining of cells with fluorescently tagged antibodies for visualization and UV-photocleavable oligo conjugated antibodies for immune cell quantification (A). Cross-sectional images of CD3^+^ T cells (pink), α-smooth muscle actin (green), and nuclei (blue) infiltration into the 150 µm and 300 µm scaffolds prototypes on explant day 14 and 30 (B; SS = silicone scaffold rung). Fold change in immune cell markers on day 14 vs. day 30 within the major pores (grey) vs. the minor pores (blue) in the 300 µm scaffold (C) or the 150 µm scaffold (D). Fold change in immune cell markers within the 300 vs. 150 µm scaffold on day 14 (grey) or day 30 (blue) within the major pores (E) or minor pores (F). Fold change in immune cell markers within major vs. minor pores on day 14 (grey) or day 30 (blue) within the 300 µm scaffold (G) or the 150 µm scaffold (H). (Significantly up- or down-regulated immune cell phenotypes are indicated by the outlined, labeled data points; *N* = 1 scaffold per explant day, *n* ≥ 9 minor or major pore ROIs; Linear mixed model with Benjamini-Hochberg multiple comparisons test for comparisons between scaffold prototypes and explant days, Multiple unpaired T test for comparisons between pore type; *p* < 0.05).

From the data, we could then characterize the impact of scaffold type, pore type, and explant day on immune-relevant protein expression using comparative volcano plots (**Figures 3C-H**). On the plots, proteins that were both significantly different (*p < 0.05*) and had a fold change > 1 are designated by a darker black outline. Markers that were significantly altered but not up-regulated by at least one fold change are summarized elsewhere (**Supplementary Table S3)**.

First, we investigated the impact of implant time (**Figures 3C-D**). For these plots, positive x-axis values represent up-regulation of proteins on explant day 30 compared to day 14. Grey points indicate the major pores, while blue points indicate the minor pores. Within the minor pores of the 300 µm rung scaffolds (**Figure 3C**), increasing the implantation time from 14 to 30 days resulted in an up-regulation in helper T cells (CD4) and their activation and targets (CD28 and MHC Class II), as well as an increase in the fibroblast marker PanCk. Within the major pores of the same scaffold prototype, the CD28 T cell activation marker was also detected, but in conjunction with the FoxP3 regulatory marker. CD163, a marker highly expressed in M2-polarized macrophages, was also down-regulated within the major pores of the 300 µm rung scaffolds over time. Within the minor pores of the 150 µm rung scaffolds (**Figure 3D**), an increased implantation time resulted in up-regulation of MHC Class II and SMA, while fibronectin and CD163 were down-regulated. For the major pores, only fibronectin and CD163 decreased over time.

Additional proteomic shifts were detected when investigating the impact of the different rung thickness geometries on host responses within the major (**Figure 3E**) or minor (**Figure 3F**) pores. Values on the positive side of the x-axis were proteins up-regulated in 150 µm rung scaffolds when compared to 300 µm rung scaffolds. Grey points indicate protein expression on day 14, and blue points indicate protein expression on day 30. Investigating the major pores, no significant alterations in protein expression were detected between the 300 and 150 µm rung scaffolds, except for a down-regulation of fibronectin in the 150 µm rung scaffold compared to the 300 µm rung scaffold at day 30 (**Figure 3E**). For the minor pores, day 14 explants exhibited no notable differences between the scaffold prototypes; however, by day 30, alterations were detected. Specifically, the 150 µm rung scaffold exhibited a down-regulation in the T cell markers CD28, Granzyme B, and CD4, the macrophage marker CD163, and the fibroblast marker fibronectin, as well as an up-regulation of SMA, when compared to the 300 µm rung scaffold (**Figure 3F**).

Finally, exploring the effect of pore type within a single scaffold geometry uncovered further alterations in host responses (**Figures 3G-H**). For these comparisons, positive values on the x-axis were proteins up-regulated in minor pores when compared to the major pores within the same scaffold prototype. Again, gray points indicate day 14 and blue points indicate day 30. For both time points, CD11c, a marker strongly associated with dendritic cells but also present on neutrophils, monocytes, macrophages, and select B cells, was significantly up-regulated within the minor pores of the 300-rung scaffolds compared to the major pores. In contrast, macrophage marker CD163 was significantly down-regulated within the minor pores on day 14 (**Figure 3G**). Examining proteins within the 150 µm rung scaffold, the minor pore region showed up-regulation of CD11c, but also showed increased detection of markers associated with myeloid-derived suppressor cells and neutrophils (Ly6G/Ly6C), monocytes (CD14) and dendritic cells (CD11c); these differences became increasingly more significant from day 14 to day 30. SMA was also up-regulated in the minor pores by day 30 (**Figure 3H**).

### Evaluation of Scaffold Geometry on Syngeneic Rat Islet Transplant Outcomes

Biocompatibility results indicated that the 3D-printed 150 µm rung scaffold prototype generated the most favorable host responses for all rung geometries tested. To evaluate its capacity to serve as a supportive environment for cellular transplantation, we chose a pancreatic islet transplantation model, as islets are particularly vulnerable to changes in their environment [64,65]. A syngeneic graft study was conducted using a diabetic *Lewis* rat model. Scaffolds fabricated using the traditional salt-leaching method served as controls, as we have previously reported their success in restoring normoglycemia using this same model [43]. Syngeneic donor islets were loaded and retained within either scaffold format using a fibrin hydrogel and transplanted into the omentum of chemically-induced diabetic *Lewis* rat recipients (**Figure 4A**).

**Figure 4:**
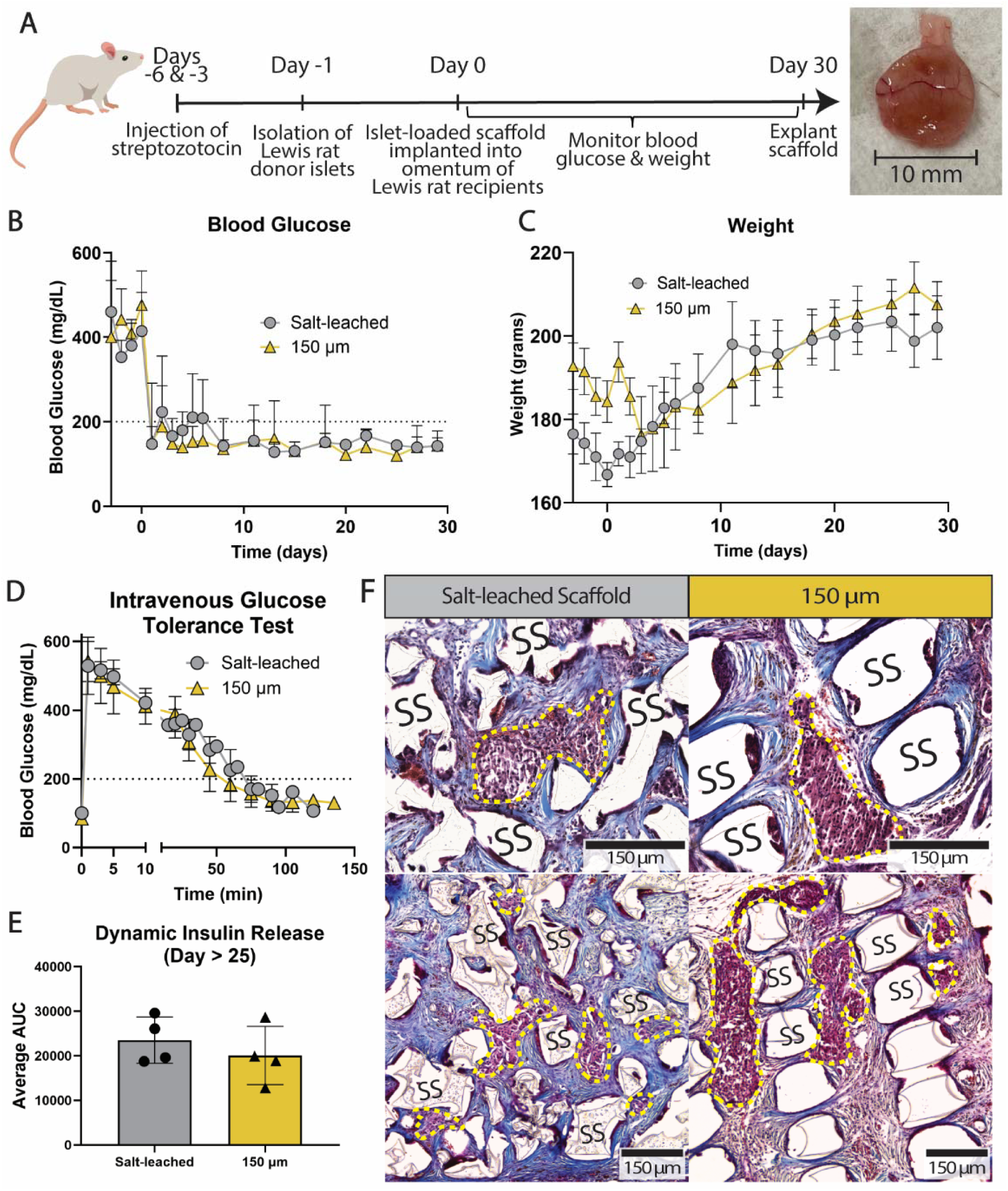
Comparison of the 150 µm 3D-printed scaffold and a traditionally fabricated, macroporous scaffold in a syngeneic *Lewis* rat islet transplant model. Timeline of islet isolation, scaffold transplantation into the omentum, and explant procedure (A). Blood glucose measurements pre- and post-scaffold transplantation (B). Weight measurements pre- and post-scaffold transplantation (C). Blood glucose measurements from an intravenous glucose tolerance test performed at least 25 days post-scaffold transplantation (D). Area under the curve measurements from the blood glucose curves of the intravenous glucose tolerance test (E). Images of Masson’s trichrome stained scaffold explants at 20x (top row) and 10x (bottom row) magnification, intra-device islets indicated by dashed yellow lines (F). (*N =* 4 rats per scaffold group; data represented as mean ± SD; Logrank Mantel-Cox test for time to normoglycemia, Two-way repeated measures ANOVA for blood glucose and weight measurements, unpaired Student’s T test for AUC measurements).

With this syngeneic model, both the 3D-printed and salt-leached scaffolds successfully and efficiently reversed diabetes in the recipients with no significant differences in the time to reversal, i.e. 1.5 ± 1 and 6.25 ± 5.377 days for the 3D-printed and salt-leached scaffolds, respectively (**Figure 4B**, *p = 0.0979*); the trend in blood glucose levels were not significant between scaffold groups (*p = 0.4663*), but time was a significant factor (*p < 0.0001*). From POD 1 to the end of the study, blood glucose was stable (*p = 0.4215*). All graft recipients gained weight over time post-transplantation (**Figure 4C**; *p < 0.0001*), but there were no significant differences between scaffold groups (*p = 0.1045*). To assess graft functionality *in vivo*, an intravenous glucose tolerance test was utilized; this test revealed no statistical differences (*p = 0.4432*) between the two groups as assessed by comparing their areas under the curve (AUC) (**Figures 4D,E**). Upon explanting the grafts, all recipients returned to their diabetic state, validating the efficacy of the transplanted islets. Histological analysis of the explanted scaffolds found viable islets distributed throughout, with high host integration and evidence of vascular infiltration for both scaffold types (**Figure 4F**).

### Scaffold Geometry Enhances Allogeneic Islet Transplantation by Accelerating Normoglycemia and Delaying Rejection

Following the encouraging outcomes of the 3D-printed scaffold within a syngeneic model, we used a more aggressive allogeneic transplant model. As recent evidence highlights the role of adaptive immune responses in FBR, we also hypothesized that an allograft transplant model may be more sensitive to alterations in scaffold biocompatibility. For this study, *Wistar Furth* donor islets were loaded within our 3D-printed 150 µm rung or salt-leached scaffolds and transplanted into the omentum of chemically-induced diabetic *Lewis* rat recipients (**Figure 5A**). This strain combination is a fully mismatched allograft that requires comprehensive immunosuppression to prevent rejection [66]. Recipients were systemically immunosuppressed using a three-drug regimen that included anti-lymphocyte serum (ALS), mycophenolic acid (MPA), and fingolimod (**Figure 5B**).

**Figure 5:**
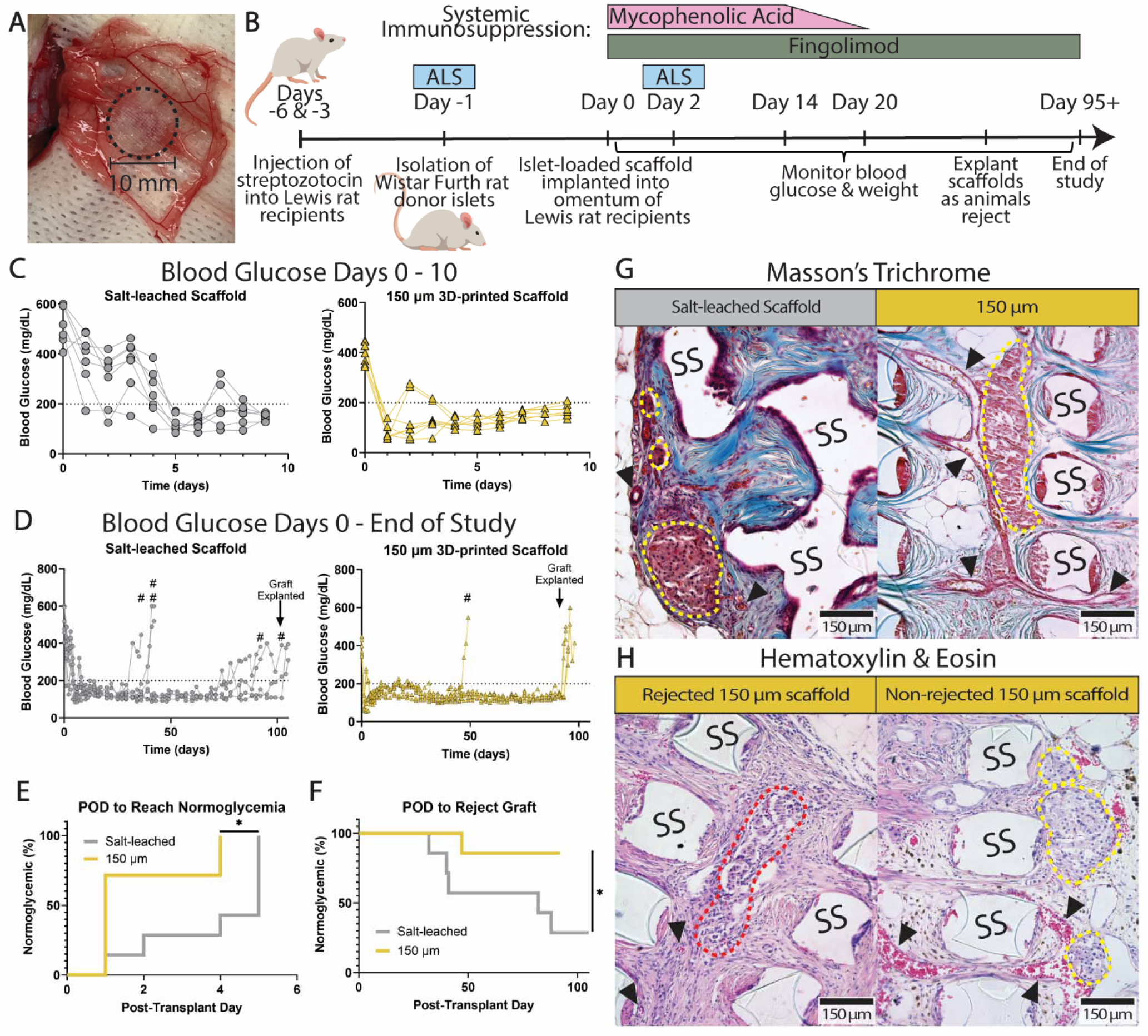
Comparison of the 150 µm 3D-printed scaffold and the traditionally fabricated, macroporous scaffold in an allogeneic rat islet transplant model. Image of an islet-loaded 150 µm 3D-printed scaffold placed on the omentum (A). Timeline of islet isolation from *Wistar Furth* rat donors, scaffold transplantation into *Lewis* rat recipients, and the systemic immunosuppressive regimen (B). Initial blood glucose readings of recipients within the first 10 days post-transplantation (C). Blood glucose readings for the entire duration of the study (D; # signifies an animal reverted back to diabetes and was sacrificed). Survival analysis curves of the time to normoglycemia (E) and time to rejection (F) for each scaffold group. Cross-sectional images of Masson’s trichrome stained scaffold explants that did not reject (G). Cross-sectional images of hematoxylin and eosin-stained 150 µm 3D-printed scaffold explants that were either rejected or not rejected (H). (SS = silicone scaffold rung, whole intra-device islets signified by yellow dashed lines, partial islet remains indicated by red dashed lines, intra-device blood vessels indicated by black carats; *N =* 7 rats per scaffold group; data shown as connected data points which signify repeated measurements on a single animal; Two-way repeated measures ANOVA for blood glucose and weight stability measurements, Logrank Mantel-Cox test for time to normoglycemia and rejection curves; \**p* < 0.05).

Blood glucose measurements were assessed up to 105 days post-transplantation (**Figure 5C,D**). Over the first 10 days post-transplant, differences in non-fasting blood glucose between the recipients of the two scaffold groups were notable (*p = 0.0005*). To investigate the time to establish normoglycemia, the 3D-printed scaffolds achieved these levels 1.86 ± 1.46 days post-transplant, while diabetic rats receiving the salt-leached scaffolds achieved normoglycemia 3.86 ± 1.68 days post-transplant. This difference was significant (*p = 0.0189*) as summarized in **Figure 5E**. From POD 11 to POD 35, the non-fasting blood glucose levels for both groups remained stable (*p = 0.3036*). However, the salt-leached scaffolds were rejected significantly faster (*p = 0.0351*) compared to the 3D-printed scaffolds (**Figure 5F**). The salt-leached scaffolds had 5 of 7 animals reject, whereas the 3D-printed scaffolds had 1 of 7 animals reject throughout the duration of the study. Of those animals, the average day to reject was POD 56.6 ± 26.2 for the salt-leached scaffold vs. POD 47 for the 3D-printed scaffold.

Animals from both scaffold groups gained weight post-transplantation (**Supplementary Figure S9**, *p < 0.0001; two-way repeated measures ANOVA*). Visualizing the scaffold explants with Masson’s trichrome revealed successful islet integration into both scaffold types. Notably, large blood vessels were observed spanning both the major and minor pores in the 3D-printed scaffolds (**Figure 5G**). Comparing rejected vs. non-rejected 3D-printed scaffold samples, we observed clear islet boundaries and blood vessel structures in the non-rejected scaffolds. Alternatively, islet fragments were observed in the rejected scaffolds with cellular infiltration within the fragments (**Figure 5H**).

## Discussion

While porous scaffolds provide critical support to cellular transplants, clinical studies have found that aggressive foreign body responses facilitate implant failure [12,15,67]. Complete fibrotic encapsulation of the implant is highly detrimental to cell survival [68–70]; however, even a modest FBR, such as sporadic fibrosis and/or the presence of inflammatory cells, can also decrease cell survival via impairment of competent vascularization and an increased exposure to cytotoxic factors [71–74]. While several parameters can modulate the severity of FBR, manipulating the device’s geometric features is easily tunable [19,38]. Most investigations into the impact of scaffold geometry on FBR have focused on pore size or overall porosity, without a clear consensus [32,35–37]. The space between the pores, i.e., the rung thickness, is a geometric feature that globally affects overall porosity; however, its impact on FBR is less understood.

To explore this phenomenon, PDMS-based porous scaffolds with distinct rung thicknesses and high biostability were created. Using a reverse-cast 3D-printing method, scaffolds with distinct pore length scales were generated. The overall fidelity of the print was robust, although the printed rungs were slightly larger and the pore size was somewhat smaller than their respective modeled sizes, likely due to PDMS swelling [46]. Also, printing accuracy decreased as the rung thickness decreased, likely due to the ∼20 µm layer resolution limitation of the 0.4 mm printing nozzle [75,76]. Overall, this printing approach yielded distinct prototypes for clearly evaluating the role of rung size on implant biocompatibility.

Following implantation, it was expected that the porous scaffold would elicit a classic host response, characterized by prompt protein adsorption, followed by neutrophil, monocyte, and macrophage infiltration, which would peak around 14 days post-transplantation [77–79]. Based on scaffold features, these infiltrating host cells would either direct a favorable or deleterious response. A healthy outcome would display features of physiological ECM deposition and vascularization. Alternatively, FBR would exhibit elevated fibroblasts with abnormally dense collagen fibers and fibronectin, resulting in fibrotic regions [18,79–84]. Further progression of the FBR would promote fibroblast to myofibroblast transition, driving capsule contraction [81–86]. The timeline for detecting this skewing to either a healthy or fibrotic response is typically one month post-transplantation.

In our study, the histological examination of the 3D-printed scaffold prototypes after 30 days *in vivo* revealed markedly different spatial responses. Classic hallmarks of FBR, e.g., dense collagen bands and tissue contraction, were detected only within the minor pores of the ≥ 200 μm rung scaffold prototypes. This response appeared localized to the minor pores, as the major pores of all implants exhibited healthy host tissue remodeling. While it is well established that device geometry influences FBR [60,87,88], this is the first report, to our knowledge, of a correlation between host FBR and rung thickness *in vivo*.

Why FBR was localized to the longer intra-pore channels is unclear, but it may be driven by host cell migratory responses. *In vitro* reports investigating the effect of channel width on cell movement found that straight and narrow channels generally promote aligned cellular migration, resulting in contraction and elongation responses [89–92]. For example, epithelial cells migrating through fibronectin-coated PDMS channels shift from a slow, spiral-like migration pattern in wider channels to a more directed contraction-elongation motion in thinner channels [89]. Similar correlations have been reported for fibroblasts and stromal cells [90,91]. Additionally, straight wound channels were found to guide cell migration parallel to the walls, resulting in actin-mediated contraction that places the cells under high tension [92]. These *in vitro* outcomes suggest that the longer pore channels, created by the thicker rungs, may hinder wound healing processes and promote a more fibrotic phenotype by altering cell migration behavior; however, other factors may also play a role.

Given the distinct differences in FBR observed between the 150 µm and 300 µm scaffolds, digital spatial profiling was employed to gain further insight into cellular features that may drive this localized response. Spatial proteomics can globally capture diverse changes within specific regions of interest of the scaffold, providing unique insight into protein alterations during host responses that are skewed towards FBR versus those exhibiting healthy remodeling.

When exploring changes in FBR-relevant innate cells and ECM markers, distinct shifts were observed. While the general macrophage marker F4/80 remained unaltered across all comparisons, a shift in macrophage phenotype, as measured via CD163, was detected. Of interest, only areas histologically exhibiting low fibrosis showed a downregulation in CD163 over time, while CD163 was unchanged within the fibrotic minor pores of the 300 μm scaffold. As such, when comparing CD163 levels within the minor pores of the two scaffold prototypes on Day 30, CD163 was decreased within the 150 μm scaffolds. While CD163 is traditionally associated with the anti-inflammatory “M2” macrophage classification, macrophage phenotype is now seen as more fluid and nuanced [93]. Furthermore, soluble CD163, generated through the proteolytic cleavage of macrophages in response to inflammatory/oxidative stimuli, is significantly increased during chronic tissue and organ fibrosis [94,95]. The ECM component, fibronectin, which plays a role in promoting the fusion of macrophages into foreign body giant cells and supporting collagen fibril formation [23,81–84], was downregulated within the 150 μm scaffold over time, as well as less abundant within both pore types of the 150 μm scaffold when compared to the 300 μm scaffold. Increased fibronectin and decreased CD163 within fibrotic regions were also reported following the implantation of microporous collagen-based scaffolds (35 - 53 μm pores), supporting our observations [96]. The other significantly altered ECM marker was SMA, which was upregulated in the minor pores of the 150 µm scaffold over time and in comparison to the larger rung scaffold on day 30. SMA may indicate either increased vascularization [15] or the presence of mature myofibroblasts and fibrosis [97,98]. When contextualized with histological observations, the increased expression of SMA is more consistent with healthy vascularization.

Other notable shifts in innate immune cells emerged when making intra-scaffold comparisons. Within the 150 μm scaffold, a notable elevation in CD11c, CD14, and Ly6G/Ly6C markers was detected in the minor pores compared to the major pores, indicating the preferential recruitment of these specific innate immune cells to this region. Classically, these markers are associated with dendritic cells (CD11c), monocytes (CD14), and neutrophils (Ly6G/Ly6C). These cell populations have been implicated in the early stage of FBR [99,100]; however, others have found these markers to associate with wound healing and resolution [98,101]. Of interest, the presence of Ly6C^+^ and CD14^+^ monocytes within scaffolds is reported to be key in driving pro-healing and anti-fibrotic responses [35]. Within the 300 µm scaffold, only CD11c was altered, with an increased presence within the minor pores. CD11c is primarily associated with dendritic cells, although monocytes can also upregulate this factor under inflammatory conditions [102]. As this marker is not broadly investigated in FBR, further investigation is necessary to classify and phenotype these CD11c^+^ cells.

While the innate arm of the immune system has long been considered the primary driver of FBR, emerging evidence highlights the significant and sometimes conflicting roles of the adaptive immune system [18,23,78]. Specific subsets of T cells, namely cytotoxic CD8^+^ T cells and helper CD4^+^ T cells, are reported to promote implant fibrosis [103,104]. Literature reports that T cells promote macrophage adhesion and fusion into foreign body giant cells [20,105]; however, the FBR timeline of these findings is conflicting, with some implicating T cells in early (< 1 month) responses, while others correlate it with advanced stages [23,105,106]. B cell involvement in FBR is also highly contradictory, with one source suggesting that B cells are critical [97] while another indicates that B cells are much less involved than T cells [103].

Alterations in adaptive immune cell markers were most prominent when comparing the minor pore regions of the two scaffold prototypes at 30 days post-implantation. Specifically, markers of helper T cells (CD4) and T cell activation and effector function (CD28 and Granzyme B) were significantly elevated within the fibrotic minor pores of the 300 μm scaffold compared to the non-fibrotic, smaller rung prototypes, indicating that activated T cells are a key contributor to the observed differences. To clearly characterize T cell phenotype impacts (e.g. Th1, Th2), further expansion of the proteomic panel is needed [17,21,23]. Of interest, increased FoxP3, a marker of regulatory T cells, was detected within the major pores of the 300 µm scaffold over time. FoxP3^+^ T cells are found to drive regenerative processes that reduce FBR, which may have contributed to the favorable host responses observed within the major pores of this prototype [104,107]. Notably, the expression of the B cell marker, CD19, remained low or below detectable levels across all comparisons, suggesting a limited role for B cells in the FBR observed in this study.

Based on the results of the initial studies, we selected the 150 μm scaffold for subsequent *in vivo* studies. This scaffold was compared to one fabricated using traditional methods (i.e. salt leaching) in both syngeneic and allogeneic rat islet transplant models. While the two scaffolds performed similarly in the syngeneic model, the impact of scaffold biocompatibility was evident in the allogeneic model, where optimized 3D-printed implants demonstrated enhanced efficacy and graft stability. Fibrosis is a hallmark of rejection in allogeneic organ transplantation and is postulated to arise from ongoing inflammation and the insufficient suppression of antigen-dependent immune responses [108–111]. Thus, it is reasonable to expect that devices that exacerbate fibrotic responses will lead to unfavorable allogeneic cellular transplant outcomes. Overall, our *in vivo* results underscore the importance of utilizing robust models to assess scaffold geometry, as well as the necessity of carefully controlling and optimizing this geometry to position the transplanted cells for successful engraftment.

While this study provides new insights into the role of geometric features in FBR, as well as the key markers involved and their impact on allograft rejection, future studies are needed to further optimize porous scaffold devices. As outlined above, a more detailed analysis of the exact phenotype of the monocytes, macrophages, and T cells involved in driving these distinct fibrotic regions could help further understand fibrotic pathways and support the identification of key therapeutic targets. Recognizing and identifying how these key adaptive immune cells modulate FBR and contribute to allogeneic rejection warrants further investigation. The optimized scaffold used herein should also be further validated within larger animal models, as FBR is typically elevated within nonhuman and human primates. Additional studies can also adapt these 3D-printed PDMS scaffolds for use in other cellular therapies beyond T1DM, such as hepatocyte or thyroid cell transplantation.

## Conclusion

In this study, a 3D-printed scaffold fabrication technique was employed in conjunction with the biostable polymer PDMS to investigate the impact of porous scaffold geometry on host FBR and islet transplant outcomes. Scaffold prototypes with distinct PDMS rung sizes ranging from 150 to 300 µm were fabricated and implanted into the epididymal fat pad of immunocompetent mice. Explant analysis revealed that larger rung sizes correlated with increased intra-device collagen deposition and adaptive immune cell infiltration, indicating that increasing the length of the pores may exacerbate the foreign body reaction to a biomaterial implant. 3D-printed scaffold prototypes with 150 µm rungs were deemed most optimal as they minimized fibrotic features. These scaffolds were compared to traditionally fabricated, salt-leached scaffolds in both syngeneic and allogeneic islet transplant models. Optimal 150 µm rung size 3D-printed scaffolds exhibited significantly superior transplant outcomes when compared to traditionally fabricated scaffolds with irregular geometry in an allogeneic transplant model. This work highlights the importance of porous scaffold geometry in minimizing foreign body reactions, thereby increasing subsequent transplant success when transitioning to more clinically relevant models.

## Acknowledgements

This work was supported by funding from Breakthrough T1D (3-SRA-2021-1033-S-B and JDRFI - 3-SRA-2017-347-M-B), and the Diabetes Research Institute Foundation (DRIF, Miami). TR Lansberry was supported by a NIH NIDDK Ruth L. Kirchstein NRSA Predoctoral Fellowship (DK138744-01A1) and the NIH NIDDK T32 Training Program in Type 1 Diabetes and Biomedical Engineering (T32 DK10876). We thank Antonello Pileggi for his consultation on the allogeneic transplant study while he was affiliated with the University of Miami. We thank Marilyn Duarte for her technical assistance during islet isolations and scaffold transplantation. We sincerely thank all members of the Stabler Laboratory and the University of Miami Diabetes Preclinical Models Core for their efforts in islet isolations, animal care, and histology. We thank the Molecular Pathology Core at the University of Florida and Kevin Johnson at the University of Miami for their assistance in processing histological samples and, specifically, Dr. Ann Dongtao Fu, for assisting with digital spatial profiling. We thank the Electron Microscopy Core at the University of Florida’s Intradisciplinary Center for Biotechnology Research (ICBR) for their assistance in using the scanning electron microscope (SEM).

## Author Contributions

TRL designed, performed, and analyzed most of the experiments, contextualized the results, and drafted the manuscript. RPA, CC, ILM, and JW contributed to experimental design, data collection, and analysis. RDM and CR contributed to experimental design and data collection for the macroporous scaffold allograft transplants and immunosuppression. CLS conceived the project, designed the research, interpreted and contextualized the data, secured funding, and contributed to drafting the manuscript. All authors contributed to the manuscript revisions.

## Disclaimer

RDM is currently employed at the US Food and Drug Administration. The work presented in this article is related to her prior employment at the University of Miami. The opinions expressed in this article are the authors’ own and do not necessarily reflect the views of the Food and Drug Administration, the Department of Health and Human Services, or the US government.

## Data availability

Data will be made available on request.

