## Supplemental Materials for "Optimizing 3D-printed Scaffold Geometry Decreases Foreign Body Response and Enhances Allogeneic Islet Transplant Outcomes"

**SUPPLEMENTARY MATERIALS**

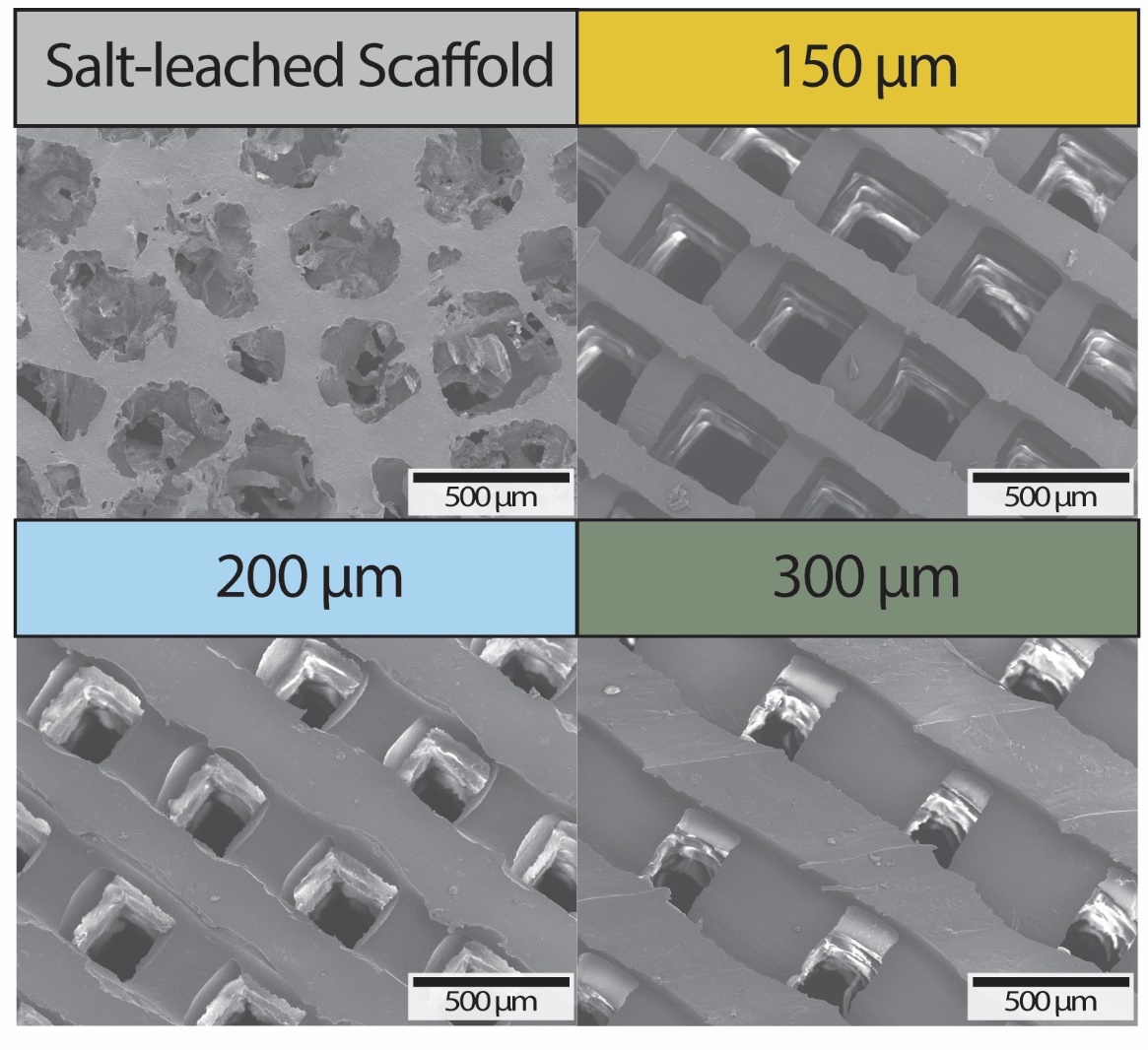

**Figure S1:** Scanning electron microscopy images of the salt-leached and 3D-printed scaffold prototypes.

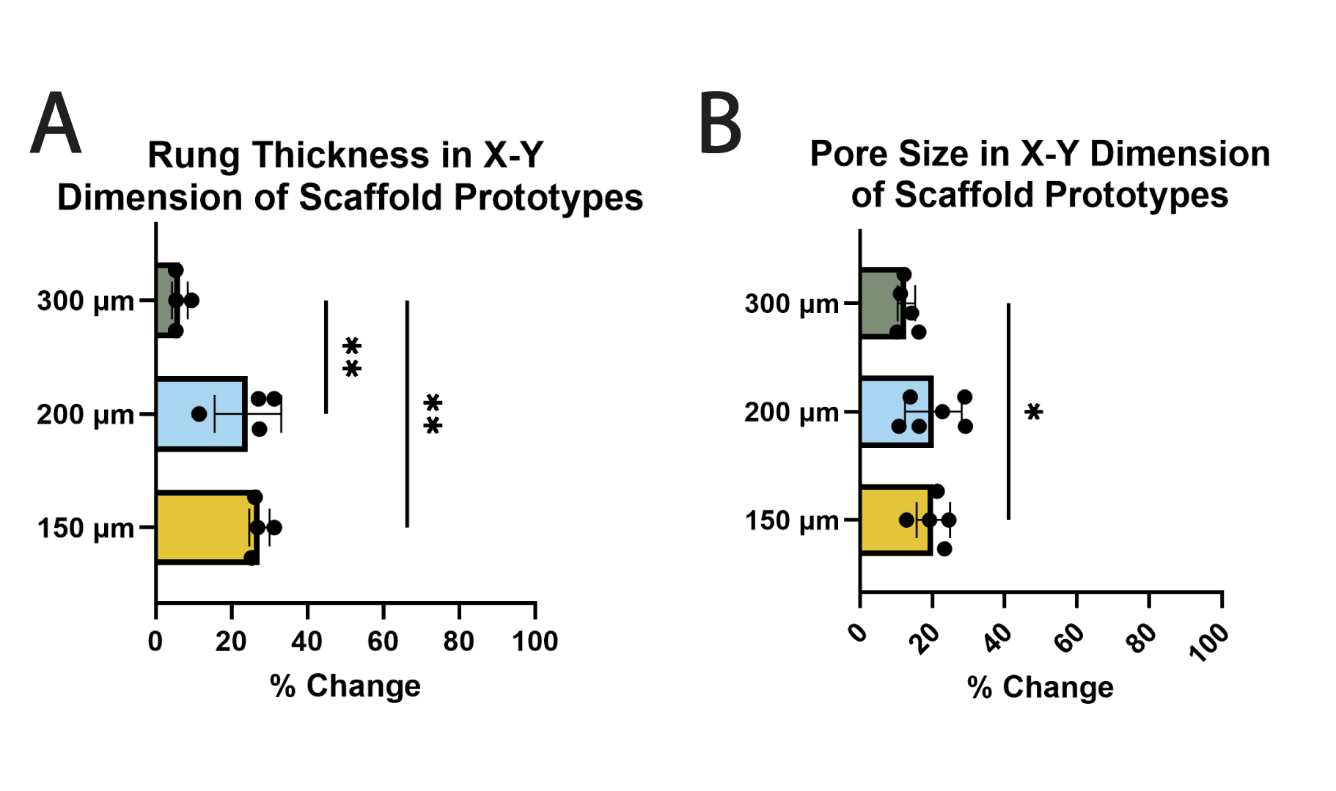

**Figure S2:** Quantification of 3D-printed scaffold fidelity from modeled to actual product. Percent change in rung thickness in the X-Y dimension from modeled to fabricated scaffold (A). Percent change in pore size in the X-Y dimension from modeled to fabricated scaffold (B). (*N* = 4 – 6 scaffolds used for all measurements, *n* ≥ 40 rungs or pores per scaffold; data represented as mean ± SD; One-way ANOVA with Tukey’s multiple comparisons test for change in rung thickness measurements, Brown-Forsythe ANOVA test with Dunnett’s T3 multiple comparisons test for change in pore size measurements; ** p < 0.05, ** p < 0.01*).

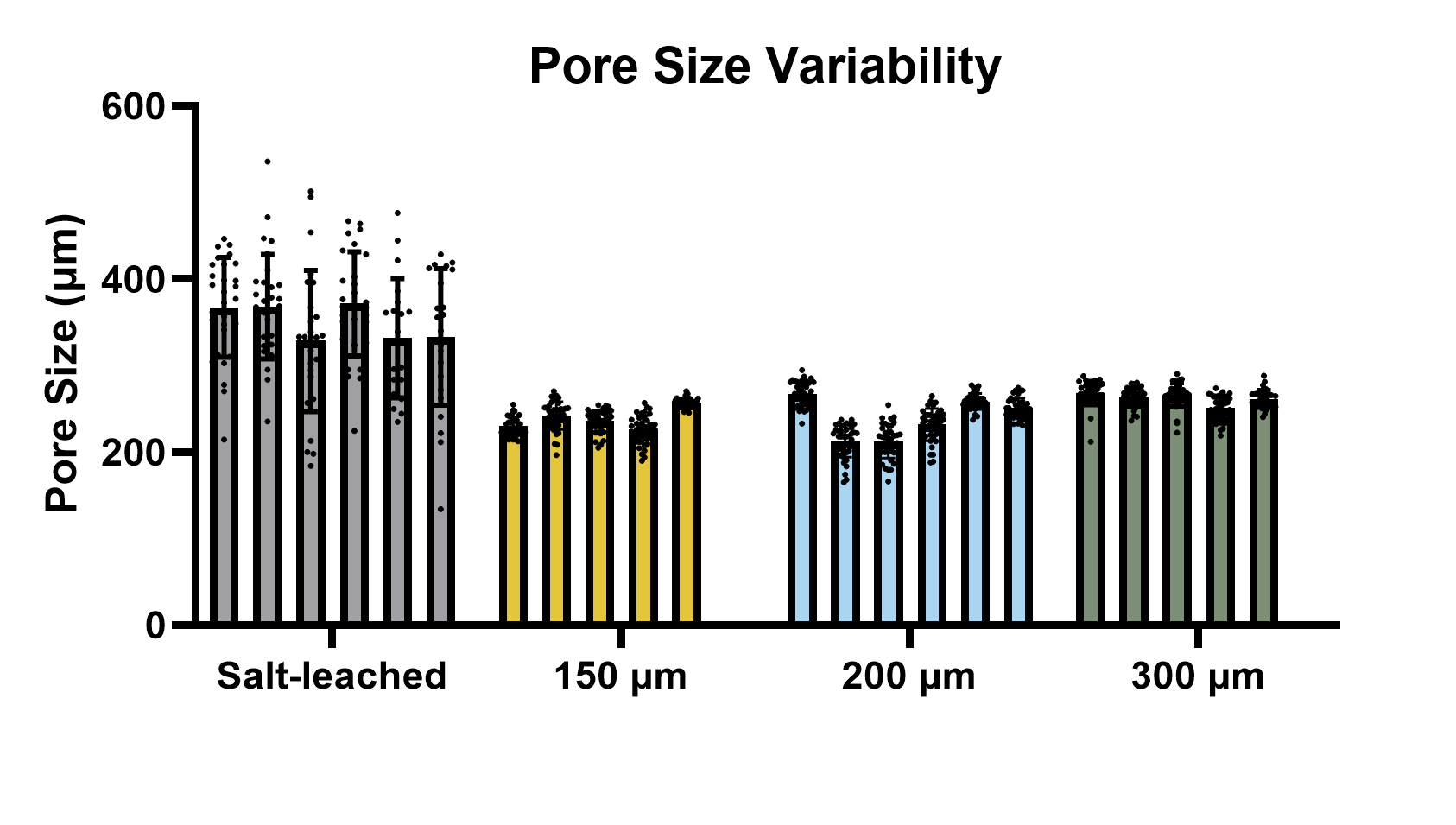

**Figure S3:** Pore size variability between the scaffold prototypes. (*N* = 5 – 6 scaffolds used per group, *n* ≥ 40 pores per scaffold; data represented as mean ± SD, each bar depicts pore size measurements for a single scaffold; Nested one-way ANOVA with Tukey’s multiple comparisons test; **** *p < 0.0001* when comparing all 3D-printed scaffold prototypes to the salt-leached scaffolds).

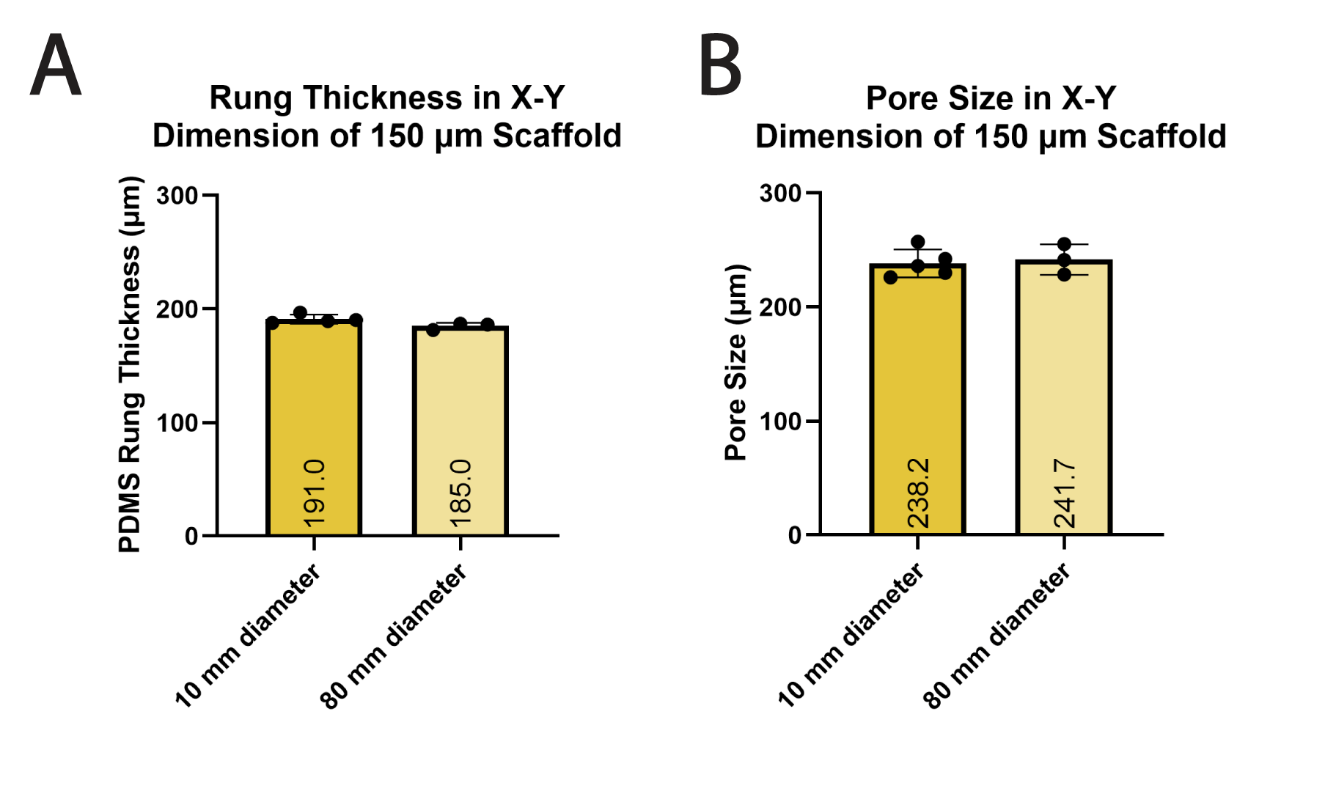

**Figure S4:** Evaluation of geometric features of scalable (80 mm diameter) scaffolds compared to their 43-fold smaller counter parts (10 mm diameter). Rung thickness measurements (A) and pore size measurements (B) of scalable scaffold compared to rodent-sized scaffold. (*N* ≥ 3 scaffolds used per group, *n* ≥ 40 rungs or pores per scaffold; data represented as mean ± SD; unpaired Student’s T test).

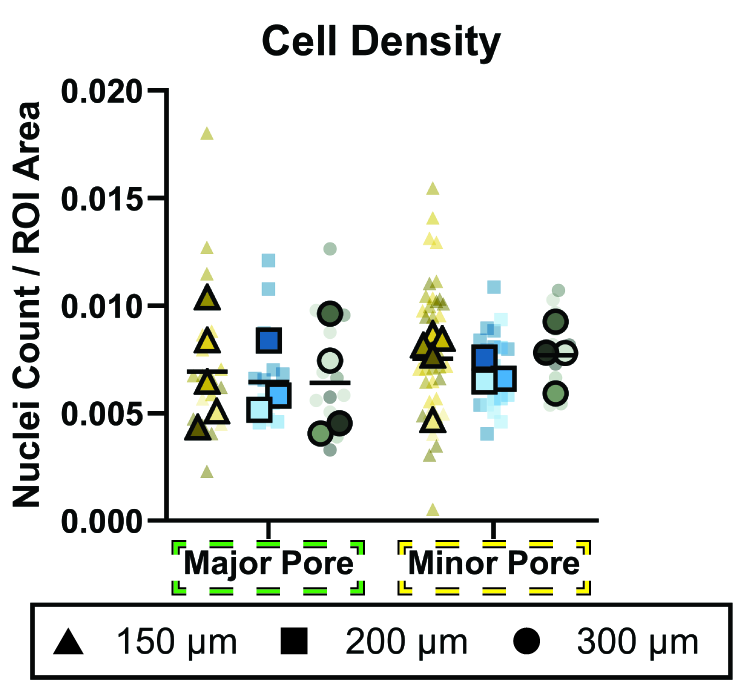

**Figure S5:** Quantification of cell density within the major and minor pores of the scaffold explants. (*N* = 3 - 5 scaffold explants, *n ≥* 3 minor pore ROIs per scaffold, *n ≥* 2 major pore ROIs per scaffold, shade of color indicates different mouse sample; all data points with mean shown; Two-way ANOVA with Tukey’s multiple comparisons test).

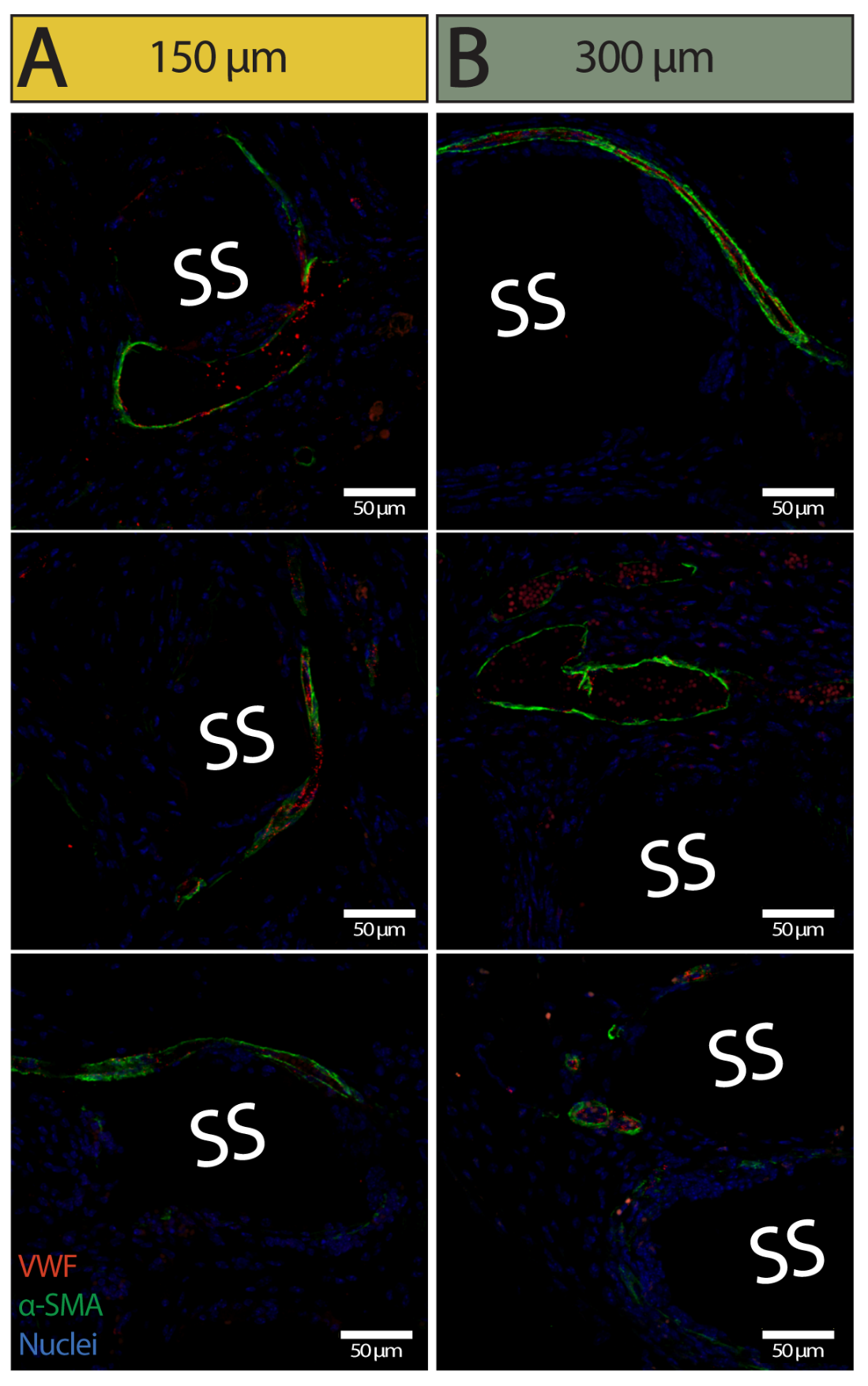

**Figure S6:** Cross-sectional immunohistochemistry images of intra-scaffold vascularization in the 150 µm (A) and 300 µm (B) 3D-printed scaffold prototypes. (VWF = von Willibrand factor, α-SMA = α smooth muscle actin, SS = silicone scaffold rung)

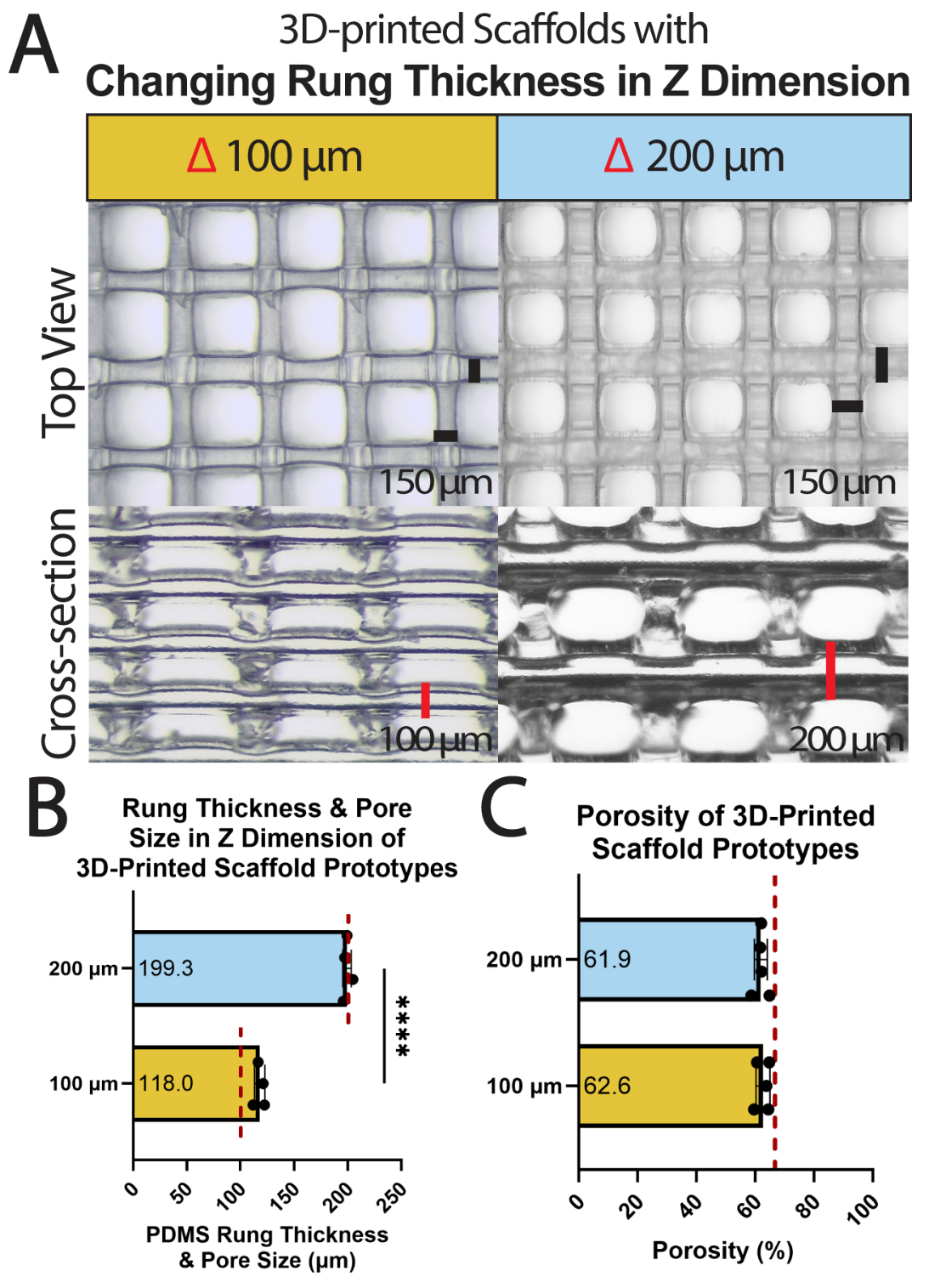

**Figure S7:** Characterization of 3D-printed scaffold prototypes with distinct PDMS rung thicknesses in the Z dimension. Top and cross-sectional brightfield images of scaffolds with 100 and 200 µm rung thickness in the Z dimension (A). Quantification of rung thickness/pore size of the scaffold prototypes in the Z dimension (B). Porosity measurements of the scaffold prototypes (C). Theoretically derived values for each parameter (red dotted line) are shown for reference. (*N* = 4 – 5 scaffolds used for all measurements, *n* = 25 rungs per scaffold; data represented as mean ± SD; unpaired Student’s T test; ***** p < 0.0001*).

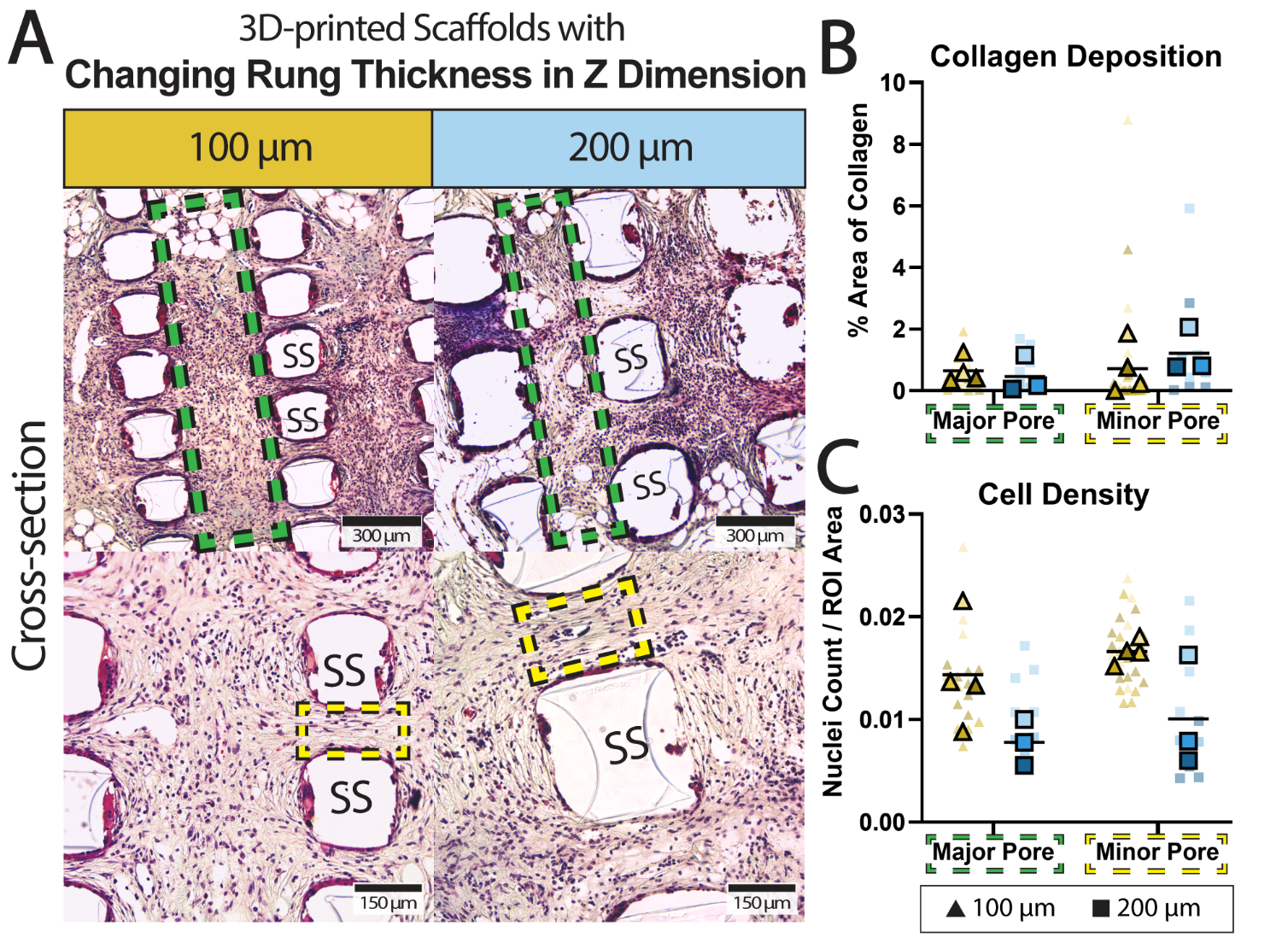

**Figure S8:** Imaging and quantification of host tissue integration into scaffold prototypes with changing rung thickness in the Z dimension. Cross-sectional images of Masson’s trichrome stained explants at 10x (top row) and 20x (bottom row) magnification (A). Quantification of collagen deposition within the major and minor pores of the scaffold explants (B). Quantification of cell density within the major and minor pores of the scaffold explants (C). (SS = silicone scaffold rung, green dashed boxes – major pore, yellow dashed boxes – minor pore; *N* = 3 - 4 scaffold explants, *n ≥* 2 minor pore ROIs per scaffold, *n ≥* 1 major pore ROIs per scaffold, shade of color indicates different mouse sample; all data points with mean shown; Two-way ANOVA with Tukey’s multiple comparisons test).

| **Protein Name** | **Annotations** |
| --- | --- |
| Rb IgG | Background |
| Rt IgG2a | Background |
| Rt IgG2b | Background |
| GAPDH | Housekeepers |
| Histone H3 | Housekeepers |
| S6 | Housekeepers |
| CD11b | DC, Myeloid |
| CD11c | DC, Myeloid |
| CD19 | B cells |
| CD3e | T cells |
| CD4 | Myeloid, T cells, Th cells |
| CD45 | Total Immune |
| CD8a | CD8 T cells, T cells |
| CTLA4 | Checkpoint, T cell Activation, T cells, Th cells |
| F4/80 | Macrophage, Myeloid |
| Fibronectin | Fibroblasts, Stroma |
| GZMB | Cytotoxicity, T cell Activation |
| Ki-67 | Proliferation |
| MHC II | Antigen Presentation, MHC2 |
| PD-1 | Checkpoint, T cell Activation, T cells |
| PD-L1 | Checkpoint, Myeloid Activation |
| PanCk | Epithelial, Tumor |
| SMA | Stroma |
| BatF3 | Antigen Presentation, DC |
| CD14 | Monocyte, Myeloid |
| CD163 | M2 Macrophage, Macrophage, Myeloid, Myeloid Suppression |
| CD28 | Checkpoint, T cell Activation, T cells |
| CD31 | Endothelial |
| CD34 | Hematopoietic |
| FOXP3 | T cells, Th cells, Tregs |
| Ly6G/Ly6C | Myeloid, Neutrophil |

**Table S1:** Immune-cell relevant protein panel for digital spatial profiling.

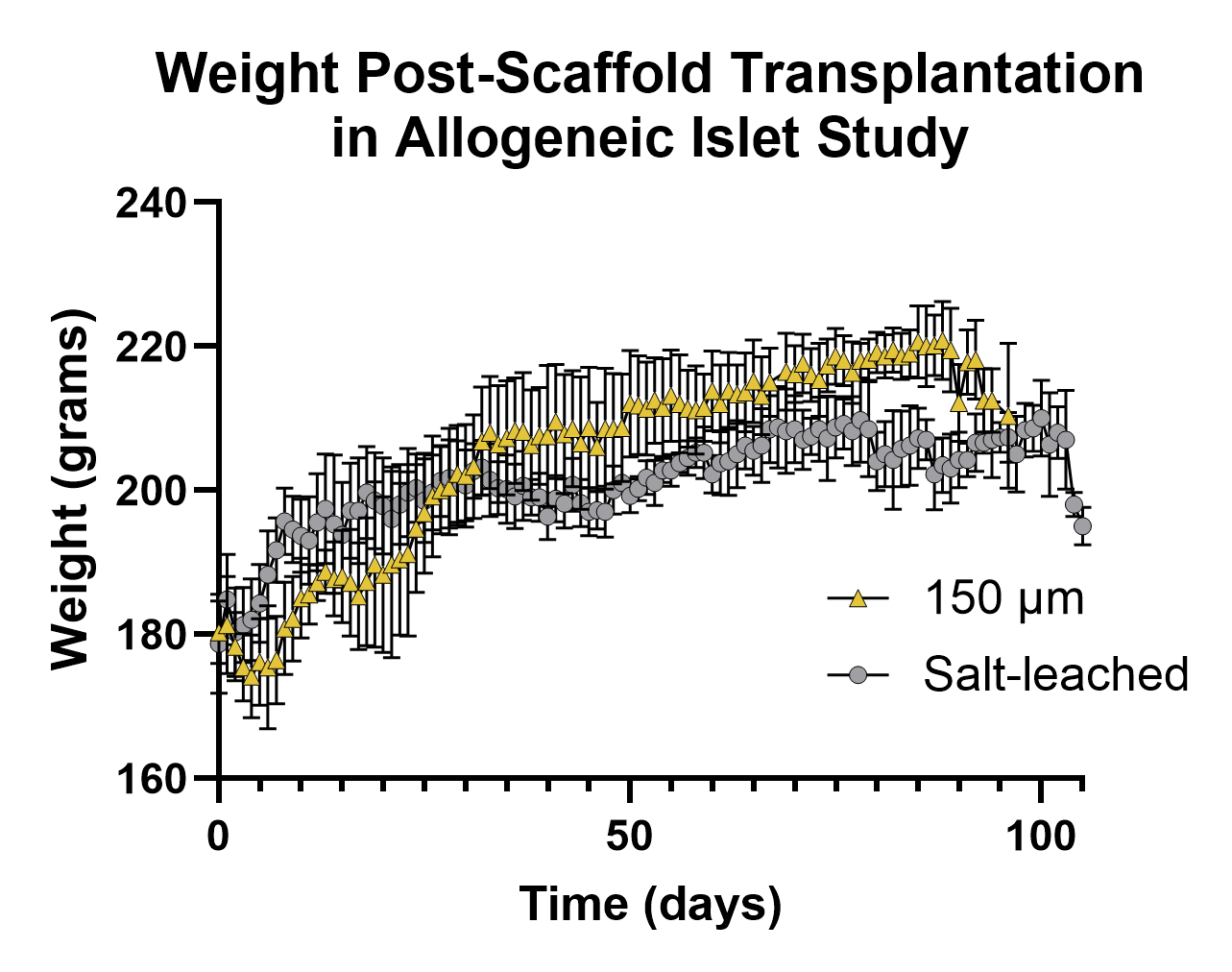

**Figure S9:** Weight measurements post-scaffold transplantation into the omentum in the allogeneic islet study. (*N* = 7 rats per scaffold group; data shown as mean ± SD; Two-way repeated measures ANOVA).

**Table S2:** Full summary of statistical analysis.

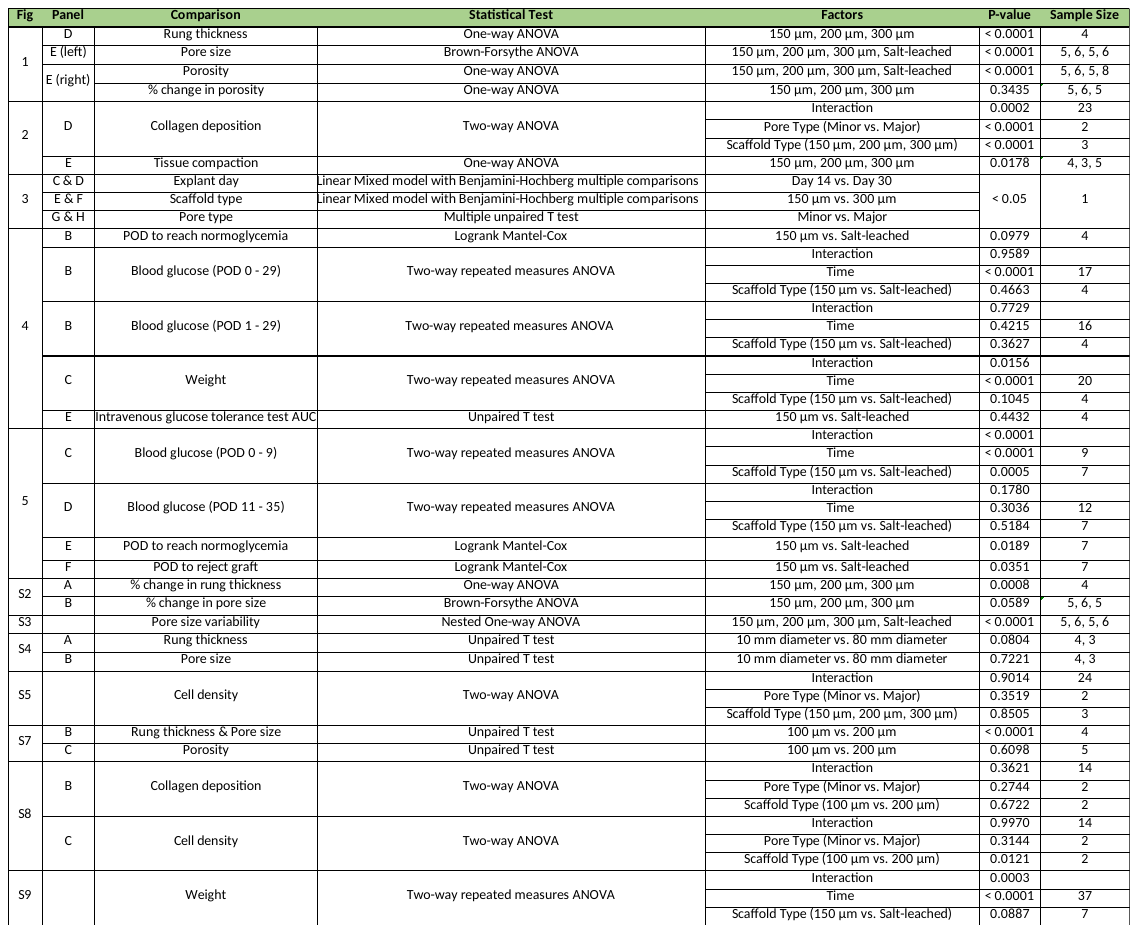
